# Interdomain linkers regulate histidine kinase activity by controlling subunit interactions

**DOI:** 10.1101/2021.08.17.456688

**Authors:** Zachary Maschmann, Siddarth Chandrasekaran, Teck Khiang Chua, Brian R. Crane

## Abstract

Bacterial chemoreceptors regulate the cytosolic multi-domain histidine kinase CheA through largely unknown mechanisms. Residue substitutions in the peptide linkers that connect the P4 kinase domain to the P3 dimerization and P5 regulatory domain affect CheA basal activity and activation. To understand the role that these linkers play in CheA activity, the P3-to-P4 linker (L3) and P4-to-P5 linker (L4) were extended and altered in variants of *Thermotoga maritima (Tm)* CheA. Flexible extensions of the L3 and L4 linkers in CheA-LVI (linker variant 1) allowed for a well-folded kinase domain that retained wild-type (WT)­like binding affinities for nucleotide and normal interactions with the receptor-coupling protein CheW. However, CheA-LV1 autophosphorylation activity registered -50-fold lower compared to WT. Neither a WT nor LV1 dimer containing a single P4 domain could autophosphorylate the P1 substrate domain. Autophosphorylation activity was rescued in variants with extended L3 and L4 linkers that favor helical structure and heptad spacing. Autophosphorylation depended on linker spacing and flexibility and not on sequence. Pulse-dipolar electron-spin resonance (ESR) measurements with spin-labeled ATP analogs indicated that CheA autophosphorylation activity inversely correlated with the proximity of the P4 domains within the dimers of the variants. Despite their separation in primary sequence and space, the L3 and L4 linkers also influence the mobility of the P1 substrate domains. In all, interactions of the P4 domains, as modulated by the L3 and L4 linkers, affect domain dynamics and autophosphorylation of CheA, thereby providing potential mechanisms for receptors to regulate the kinase.

## Introduction

Chemotactic bacteria coordinate their motility to changing environmental conditions by sensing specific chemicals through a receptor-mediated process^1-4^. Increasingly, chemotaxis is being recognized as a key determinant of pathogenicity in motile bacteria and serves as a paradigm for signal transduction systems in general^5-10^. A highly conserved supramolecular protein infrastructure transduces chemoreception signals to regulate the motility engines of the cell^3^. Dimeric transmembrane chemoreceptors, also known as methyl-accepting chemotaxis proteins (MCPs), bind attractants and repellants in the periplasmic space and undergo conformational shifts that cross the cell membrane and reach the cytosolically distal end of the receptor dimers, where the signal-transducing histidine kinase CheA binds^3,4,10,11^. Sensitivity to small changes in ligand concentration over a large dynamic range permit chemotactic bacteria to track chemical gradients effectively^24^. The core chemosensory components form extended hexagonal arrays consisting of trimers of receptor dimers (TODs), the CheA kinase, and an adaptor protein CheW^2^·^11^-^12^. The array superstructure facilitates cooperative responses and signal gain. Receptor conformational signals modulate CheA autophosphorylation and therefore subsequent phosphoryl-transfer to the response regulator CheY. Phosphorylated CheY (CheY-P) binds to the flagellar rotor and directly affects flagellar rotation^12^. The molecular details of CheA autophosphorylation are therefore of central importance to understanding signal processing and transduction in chemotaxis^4^.

CheA is comprised of five domains: a P1 substrate domain, a P2 docking domain, a P3 dimerization domain, a P4 kinase domain, and a P5 regulatory domain that is homologous to CheW^13^. Linking residues that reside outside of the respective domain secondary structures connect the five units. The P1-to-P2 and P2-to-P3 linkers (hereafter referred to as L1 and L2, respectively) are long and flexible (72 and 38 residues, respectively for *Tm* CheA), whereas the P3-to-P4 and P4-to-P5 linkers (hereafter referred to as L3 and L4, respectively) are short (2-3 amino acids each) **(Figure 1A)**^13-16^. The catalytic cycle of CheA involves P4 binding to ATP, subsequent binding of P4 to the P1 domain of the adjacent subunit, transfer of the y-phosphate to a conserved histidine residue on P1, and P1 release^14,17-22^. ADP then exchanges for ATP in the P4 active site for the next round of catalysis following phosphotransfer from P1 to a conserved CheY aspartate residue^23^. Within the arrays, the P5 domain of CheA and its paralog CheW form rings that assemble the signaling units into extended hexagonal arrangements **(Figure 1B)**^24-27^. Conformational signals affecting CheA propagate vertically along receptors and laterally throughout the arrays^1-4^.

**Figure 1.**
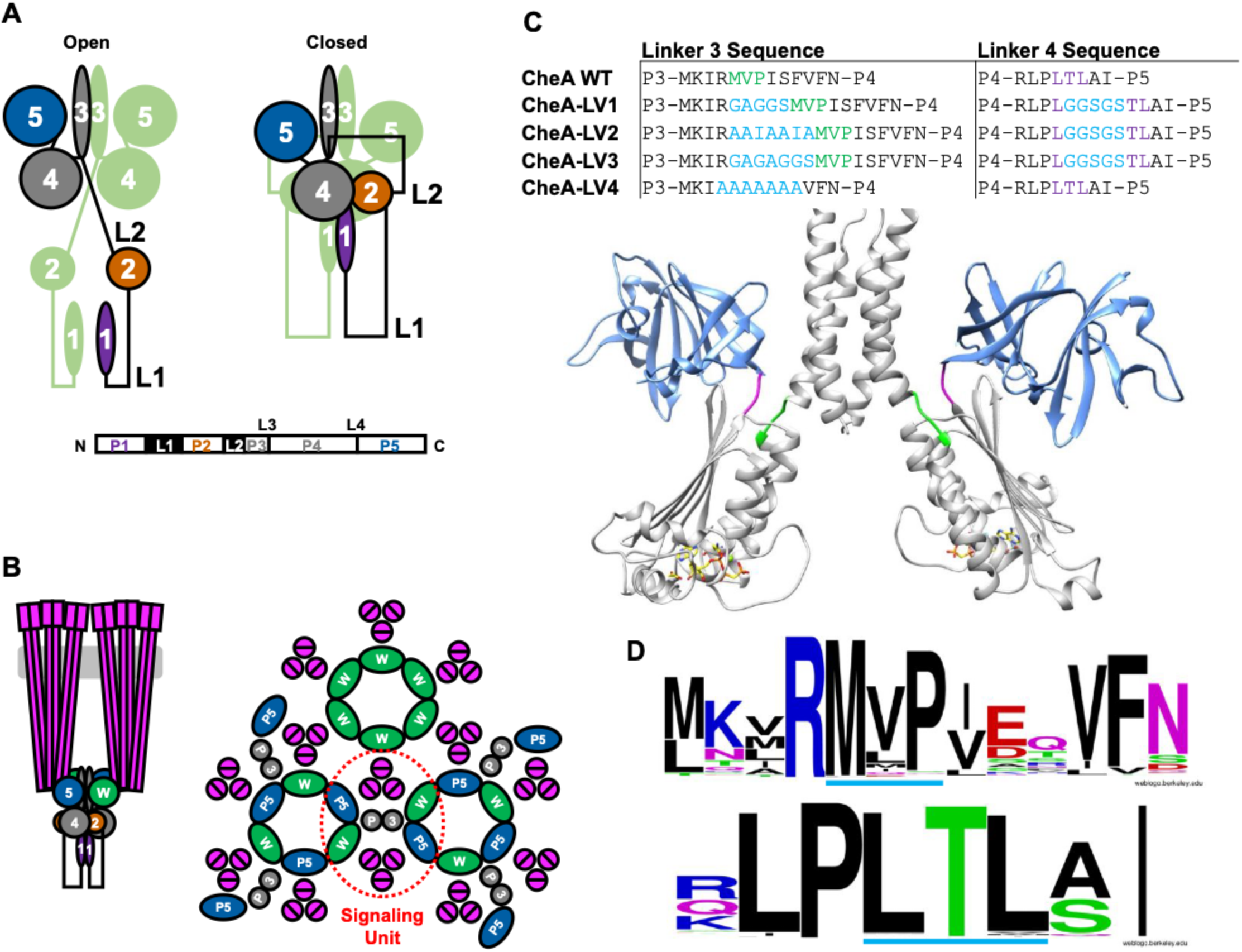
Domain arrangements, linker conformation and linker sequence conservation of CheA. **(A)** The domain arrangements of *T. marìtima* CheA WT in open and closed conformations. Schematic of the closed state is based published crystal structures of domains and complexes (P1 from PDB ID: 1TQG; P2 from PDB ID: 1UOS; P3P4P5 from PDB ID: 1B3Q) and other biophysical data that includes interdomain crosslinking; the open state is less well characterized, although it involves a greater mobility of the P1 and P2 domains^28^’^31^. The P1 and P2 domains are connected by a long flexible linker (L1) and the P2 and P3 domains are connected by another long flexible linker (L2). Relatively short L3 and L4 linkers connect the P3 domain to the P4 domain and the P4 domain to the P5 domain, respectively. **(B)** The chemotaxis receptor;kinase array architecture. Left: Vertical view of transmembrane Trimers of MCP Receptor Dimers (TODs) that contact CheA and CheW (green) in the cytoplasm and regulate CheA autophosphorylation. Right: a lateral view of the extended arrays formed by neighboring signaling units. Two TODs, 2 CheW subunits, and a CheA dimer constituting a core signaling unit are circled in red. (C) Above: table showing linker modifications of CheA variants. The WT L3 and L4 residues are colored green and magenta, respectively. Residue insertions or changes are colored in cyan. Linker variants (LV) 1,2, and 3 extend the L3 and L4 linkers. LV1 and LV3 are designed to maximize flexibility. The LV2 sequence inserts a heptad repeat that favors α-helical structure. LV3 would also allow an in-register α-helix but maintains greater flexibility than LV2. LV4 has a WT L4 length, but a poly-Ala sequence. Below: model of dimeric TmCheA P3P4P5 from PDB ID: IB3Q highlighting L3 and L4 (P3P4:gray, P5:blue, L3:green, L4:magenta, ADPCP shown as a stick model). **(D)** Sequence logos of the CheA L3 (top) and L4 (bottom) elements across 501 CheA sequences that include those from *Thermotoga maritima, Escherichia coli, Salmonella enterica, Helicobacter pylori, Bacillus subtilis, Borrelia burgdorferi, Treponema pallidum,* and *Campylobacter jejuni.* The L3 and L4 linker positions are underlined in light blue for clarity.

In general CheA kinases display a basal level of autophosphorylation, which can be increased or decreased upon interaction with chemoreceptors, depending on their state of ligand occupancy and the particular CheA^28^. Thermostable *Thermotoga maritima (Tm)* and *E. coli* CheA have been described to assume a receptor-complexed “closed” conformation that constrains the P1 domains and an “open” conformation that enables autophosphorylation and phosphotransfer **(Figure 1A)**^4,28-31^. P1 phosphorylation decreases dramatically upon inhibition by receptors^29^. The molecular details of the kinase conformations responsible for different autophosphorylation levels are not fully characterized for any CheA. The isolated P4 domain phosphorylates free P1 domain with very low activity compared to full-length CheA^4,30^. However, for *Tm* CheA, fragments composed of only the P3 and P4 domains increase kinase activity above full-length CheA (CheA FL) and well above P4 alone. Dimerization conferred by P3 increases kinase activity perhaps because the L3 connection affects P4 activity either by direct influence over the P4 catalytic machinery or by controlling the proximity of the P4 domains in a CheA dimer^4,30^. The P4 domains are closely associated with P1 and P2 in the inhibited state of *Tm* CheA^31-33^ and in *E. coli* arrays, greater order is observed in the regions of the P4 domains when the kinase is bound to inhibitory receptors^34^. The release of constrained P1 from the kinase core domains (P3-P5) when P4 binds non-hydrolyzable ATP analogs increases the hydrodynamic radius and flexibility of the CheA dimer^30,31^. Hence, the state of the P4 domain influences the interactions of the P1 substrate domains. Nonetheless, covalent connection of the P1 domains is not essential for activity. For *E. coli* CheA, free P1 domains can be phosphorylated by receptor-bound P3-P4-P5 units and the activity level responds to receptor occupancy^35^. Inhibition of CheA may involve exclusion of P1 from the P4 ATP binding site by regulating the accessibility of either module^4,24,36^.

Cryo-electron tomography of CheA arrays coupled with molecular dynamics (MD) simulations suggest that the position, dynamics, and hydrogen-bonding network of the P4 domain may be coupled to a secondary structure change in the L3 linker^37^. Additionally, distinct conformations of the P4 domains in "dipped" and "undipped" conformations support the notion that regulation of P4 mobility is important in the CheA catalytic cycle^26,36,37^. Observations that increased proximity between the P4 domains is associated with the inhibited state of the kinase in pulse-dipolar electron-spin resonance (ESR) experiments corroborate this hypothesis^31^. Importantly, changes to the L3 and L4 linkers have been demonstrated to affect P4 kinase activity; point mutations of the L3 residues modestly reduce the basal autophosphorylation level of CheA, but also impair kinase activation by receptors^38,39^. Residue substitutions in the L4 linker, in contrast, both decrease and increase the basal autophosphorylation level of CheA even in the absence of the P5 domain and also impair the ability of chemoreceptors to modulate CheA activity^38,40^. In addition, strong negative cooperativity in nucleotide binding between the subunits of dimeric CheA further suggests that the P4 domains influence one another, either through direct contact or through conformational propagation. Restriction of P4 movements by the L3 and L4 linkers, therefore, may play a role in regulating kinase activity^41^.

To investigate the influence of the L3 and L4 linkers on CheA activity, we have produced and characterized a set of *Tm* CheA variants wherein the L3 and L4 linkers are altered and extended to increase their flexibility, favor helical conformational states and alter P4 juxtaposition **(Figure 1C)**. These modifications serve to disconnect the P4 domain from the P3 and P5 domains and enforce alternate physical linkages. The linker variations produce large changes in CheA activity despite the proteins maintaining near normal interactions with CheW and receptors. Subunit exchange experiments coupled with phosphorylation assays, small-angle x-ray scattering (SAXS), cross-linking data, and temperature dependencies of autophosphorylation imply that the basal level of kinase activity is tightly coupled to the conformational properties and interactions of the P4 domain, as influenced by the L3 linker. The co-dependencies of these structural features provide a possible conduit for conformational signals from the receptors to regulate CheA.

## Methods

### Chemicals, Reagents, and Proteins

All proteins used in the experiments presented in this study were derived from *Thermotoga maritima* and produced in *E. coli.* CheY (UniProt ID: Q56312), CheW (UniProt ID: Q56311), CheA WT (UniProt ID: Q56310), CheA P1-3 (residues 1-354), CheA-LVI, CheA-LV2, CheA-LV3, CheA-LV4, and *Escherichia* co//Tar Foldons Short (Tar_FO_ Short) were recombinantly expressed in BL21 *E coli* cells using a pet28a vector and purified by affinity chromatography as previously reported^29^. 2’(3’)-O-(2,4,6-trinitrophenyl)-adenosine 5’-triphosphate (TNP-ATP) was purchased from Invitrogen as a trisodium salt and stored in the dark at -20 °C at 10 mg/ml_. DSSO was purchased from ThermoFisher. Copper (II) sulfate and (1,10)-phenanthroline were purchased from VWR and subsidiaries. Tris, HEPES, KCI, NaCI, MgCI_2_ were purchased from Sigma. [γ-^32^P]ATP was purchased from Perkin Elmer. ADP-NO was synthesized as previously reported^42^.

### Rosetta Loop-Modeling, Rosetta Relax with RosettaEPR constraint, and RosettaRemodel

Peptide linkers of various sequences were inserted into the L3 and L4 of CheA models that consist of P3, P4 and P5 domains. The CheA P4 domain was adjusted in position to accommodate the inserted designed linkers in L3 and L4. Rosetta loop modeling was then applied using the Kinematic Closure (KIC) method to calculate allowable torsion angles of the backbone, which generate loop conformations to close the breaks in the protein chain^43,44^. 250 KIC build attempts were made with 500 models generated. The best model for each linker variant was selected based on the score in Rosetta Energy Units (REUs) and finalized after structural comparison with the other top models.

These top-scoring CheA models of LV1-4 were selected to run in the Rosetta Relax program, an all-atom refinement by minimization and sidechain packing in the Rosetta force-field^45-17^. The PDS DEER (Double Electron-Electron Resonance)-derived ADP-NO distances (see below) between the ATP-binding pockets of a CheA dimer were used to provide an additional constraint in relaxing the structure^48^. To maintain these constraints, a suitable function based on the “motion-on-a-cone” model developed by Alexander *et al.* (2008) was implemented in RosettaEPR^48,49^. The function describes the relationship between the experimental measured spin label distance (dSL) and Cβ-Cβ distance (dCβ) in which an acceptable range of dCβ is determined from dSL (dCβ e [dSL-12.5 Â, dSL +2.5Â]). The output models were then evaluated based on their Rosetta all-atom score for analysis. In the Rosetta Relax productive run, 100 models of each CheA variant were generated with five cycles of minimization and sidechain repacking^45^. Top models based on the REU scores were selected and analyzed.

RosettaRemodel calculates the backbone design of a protein model using structural fragments based on secondary structure assignments and stereochemical parameters from entries of the PDB^50^. In this approach, secondary structures can be inserted into a protein model by assigning fragment types such as Ή", "L", Έ" or “D”, which stands for helix, loop, extended strand or degenerate/random, respectively. In this case, “D” was chosen to assign H/L/E randomly to the designed linker residues to avoid secondary structure bias. The CheA models of LV1-4 produced from RosettaRemodel were run in the Rosetta Relax program with RosettaEPR, using the aforementioned PDS-derived distance constraints^48,50^. 100 models were generated with five cycles of minimization and sidechain repacking.

### CheA Autophosphorylation Assays

CheA autophosphorylation was monitored by ^32^P incorporation. All radioassays were carried out in 50 mM MOPS pH 7.5, 150 mM KCI, 10 mM MgCI_2_. CheA WT and CheA LV samples were prepared at between 1.5-3 µM protein (consistent within each experiment) at a total volume of 23 µL. Following room temperature incubation for -15 minutes, [γ-^32^P]ATP was added to a final concentration of 1 mM and the reaction was quenched after 12 minutes using 4x SDS-PAGE loading buffer containing 50 mM EDTA pH 8.0. Elevated temperature experiments employed a heating block. The samples were loaded onto a 4-20% Tris-glycine polyacrylamide protein gel purchased from Invitrogen. Gel electrophoresis was carried out for 35 minutes at 125 V constant voltage, and gels were dried in a Bio-Rad Gel Dryer overnight in cellophane. Exposure to the storage phosphor plate occurred for >20 hrs prior to imaging on a Typhoon Image Scanner. Gel bands were quantified using ImageJ. Some gels were rehydrated in water and stained using 10% Acetic Acid, 25% Ethanol in water containing 2.5 g/L Coomassie Brilliant Blue, and subsequently destained with 10% acetic acid and 25% ethanol.

### CheA Subunit Exchange and Disulfide Crosslinking

CheA WT, CheA-LVI, and CheA P1-3 variants containing single cysteine residues at only positions 45 or 492 were reconstituted in 50 mM MOPS pH 7.5, 150 mM KCI, 0.5 mM MgCI_2_, 5% glycerol at 20 µM and following either brief incubation at room temperature or a 5 minute incubation at 55 °C to exchange subunits, Cu(ll) and (1,10)-phenanthroline were added to final concentrations of 5 and 15 mM, respectively. The reactions were quenched with loading buffer after 8 hrs of incubation at room temperature^51^. The resulting samples were subjected to gel electrophoresis on a 4-20% Tris-glycine protein gel. The gels were stained in 10% acetic acid, 25% ethanol in water containing 2.5 g/L Coomassie Brilliant Blue and destained in 10% acetic acid, 25% ethanol solution. The gel was photographed and the lanes quantified using ImageJ.

### CheA CheW Crosslinking

Samples of 10 µM CheA with and without 10 µM CheW were incubated in 50 mM HEPES pH 8.0, 150 mM KCI, 0.5 mM EDTA, 10 mM MgCI_2_ and crosslinked with 12.7 mM disuccinimidyl sulfoxide (DSSO) for 40 minutes^52^. The reaction was quenched with 20 mM Tris buffer, and the samples subjected to gel electrophoresis on a 4-20% Tris-glycine protein gel. The gel was stained in 10% acetic acid, 25% ethanol in water containing 2.5 g/L Coomassie Brilliant Blue and destained in 10% acetic acid, 25% ethanol solution.

### TNP-ATP Binding

All experiments were performed in 50 mM Tris pH 8.0, 150 KCI, 10 mM MgCI_2_ at room temperature in a quartz cuvette with continual stirring via a magnetic stir bar. Aliquots of TNP-ATP were added to a temperature-controlled quartz cuvette containing 1 mL of CheA (1 µM subunit, 0.5 µM dimer). Binding isotherms at 20 °C were constructed using integrated fluorescence between 500 and 650 nm following excitation at 410 nm. Integration was performed using the trapezoid rule (average of left and right Riemann sums). Effective IC_50TNP_. atp values were calculated using MATLAB by fitting the isotherm to the following equation, where *f_TNP_. _ATP_(bound)* indicates fractional fluorescence:

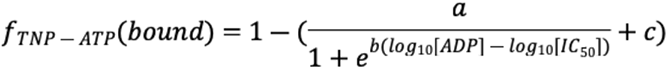

To calculate effective IC_50_ values for ADP, CheA variants and TNP-ATP were added to solution in equimolar amounts (5 µM) in a 1 cm x 1 cm fluorescence cuvette and binding was monitored by fluorescence emission at 550 nm with an excitation wavelength of 410 nm^53^. ADP was added from a stock concentration of 1 M. Effective IC_5_oadp values were calculated using MATLAB:

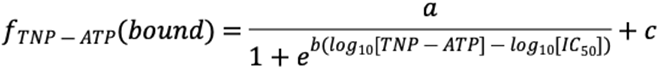

### TNP-ATP/ADP Chase Experiment

CheA WT (5 µM) and TNP-ATP (5 µM) were incubated at room temperature and stirred continuously with a magnetic stir bar. The sample was excited with 410 nm light and fluorescence emission at 550 nm was monitored. 5 µL of 1 M ADP was added and the fluorescence decay monitored. Fluorescence decay was fit to an exponential decay function using MATLAB. CheA V350C was used out of convenience. CheA V350C does not differ from CheA WT in its nucleotide binding properties or activity profile.

### Size-Exclusion Chromatography Small-Angle X-Ray Scattering (SEC-SAXS)

SAXS data were collected at CHESS G1 line on a Finger Lakes CCD detector. 100 µL of CheA WT and CheA-LV1 samples containing 4.5-5.5 mg/mL protein was injected onto a Superdex 200 Increase 10/300 Column pre-equilibrated with sample buffer (50 mM HEPES pH 8.0, 150 mM KCI, 10 mM MgCI_2_) coupled to the G1 SAXS sample cell. The monodisperse stream was fed through a Wyatt MALS/DLS and dRI detector and data was collected at 4-10 °C with 2 second exposure times to the x-ray beam at a flow rate of roughly 0.2 ml/min. RAW was used to process the SEC-SAXS data and to generate Guinier Fits and dimensionless Kratky Plots^54,55^. FoXS was used to fit the scattering data to a structure of the full-length CheA dimer in its inhibited conformation and compare experimental scattering profiles to the expected scattering profile projected from the model^56,57^.

### ESR Spectroscopy Measurements

Proteins were incubated with more than a 500-fold molar excess of ADP-NO at 4 °C for upwards of 12 hrs. Continuous wave ESR (cwESR) experiments were carried out at X-band (-9.4 GHz) on a Bruker E500 spectrometer. The cwESR experiments were performed at room temperature with a modulation amplitude of 1-2 G. For 4-pulse-DEER, the samples were exchanged into deuterated buffer containing 30% d®-glycerol, checked by cwESR and plunge frozen into liquid N_2_. The pulse ESR measurements were carried out at Q-band (-34 GHz) on a Bruker E580 spectrometer equipped with a 10 W solid state amplifier (150W equivalent TWTA) and an arbitrary waveform generator (AWG). DEER was carried out using four pulses (π/2-Tļ-π-Tļ-ττ_P_ump-T2-π-τ_2_-echo) with 16-step phase cycling at 60 K. The pump and probe pulses were separated by 56 MHz (-20 G). The signal background was approximated by a polynomial function in the semi-log scale and subtracted out. Noise from the time domain data was then removed by the WavPDS method (a wavelet denoising procedure for PDS) and distance distributions of spin separations calculated by the singular-value decomposition (SVD) method, developed to solve ill-posed problems such as for the PDS signal and estimate uncertainty in their measurement^58,59^.

## Results

### Design of the CheA Linker Variants and their Purification

Despite being far removed from the catalytic machinery, residue substitutions within the L3 and L4 linkers alter CheA basal activity, dynamics, and activation. Thus, we decided to test the effect of specific structures and conformational properties in these regions on autophosphorylaton activity of the *Tm* CheA, for which there is extensive structural data, and we have established methods for structural modeling^4,31^. In LV1 we chose L3 and L4 linkers that extend and detach the P4 domains from the P3 domains by flexible regions yet leave the ATP-binding and autophosphorylation machinery largely undisturbed. LV2 contains a heptad spacing and a sequence that favors helical propensity to recover the relative orientation of the P4 domains found in the WT. LV3 maintains the heptad spacing of LV2 but adopts the linker flexibility of LV1. To evaluate the importance of specific sequence, LV4 retains the WT spacing but converts the linkers to poly-alanine. Modification of the L4 linker was limited to flexible extension in LV1-3. The P4 domain was defined in full-length *Tm* CheA as residues 355-538 based on crystal structures of the P3-P4 unit (PDB code: 4XIV) and of the P4-P5 unit (PDB code: 2CH4)^30,60^·^61^. The amino acid sequence of all CheA variants (LV1, 2, 3, 4) are identical to that of CheA WT save for the alterations to L3 and L4. To assess whether the linkers conform to the prescribed properties, each conformation was modeled extensively in Rosetta using unbiased loop building and energy minimization. RosettaRemodel was used to generate random secondary structural elements for the fixed L3 sequence, which were then fed into the RosettaRelax program and subjected to five rounds of minimization and sidechain repacking^45,50^. Models with top scores in Rosetta Energy Units (REUs) were selected for comparison of L3 across CheA variants **(Figure 2AB).** As expected, LV1 formed an irregular loop that contained little secondary structure, LV2 was α-helical, as was LV3, although LV3 had less helical propensity as judged by reduced adherence to probable Ramachandran stereochemical values, which are favored in α-helices **(Figure 2B)**. The increased L3 helicity upon addition of two residues in LV2 and LV3 likely results from the linker spacing allowing continuous helix formation between P3 and P4 while maintaining their relative positions. In LV4, L3 formed a partially helical loop that closely mirrored the L3 of CheA WT. CheA WT and all LV proteins expressed well in *E. coli* cells using standard methods. Size-exclusion chromatography (SEC) was used to isolate the dimeric fractions of each preparation and small-angle x-ray scattering (SAXS) coupled to SEC was used to confirm oligomeric state and monodispersity **(Figure S1)**. In these respects, the purified WT and linker variants behave similarly to past samples of full-length *Tm* CheA^30,31^.

**Figure 2.**
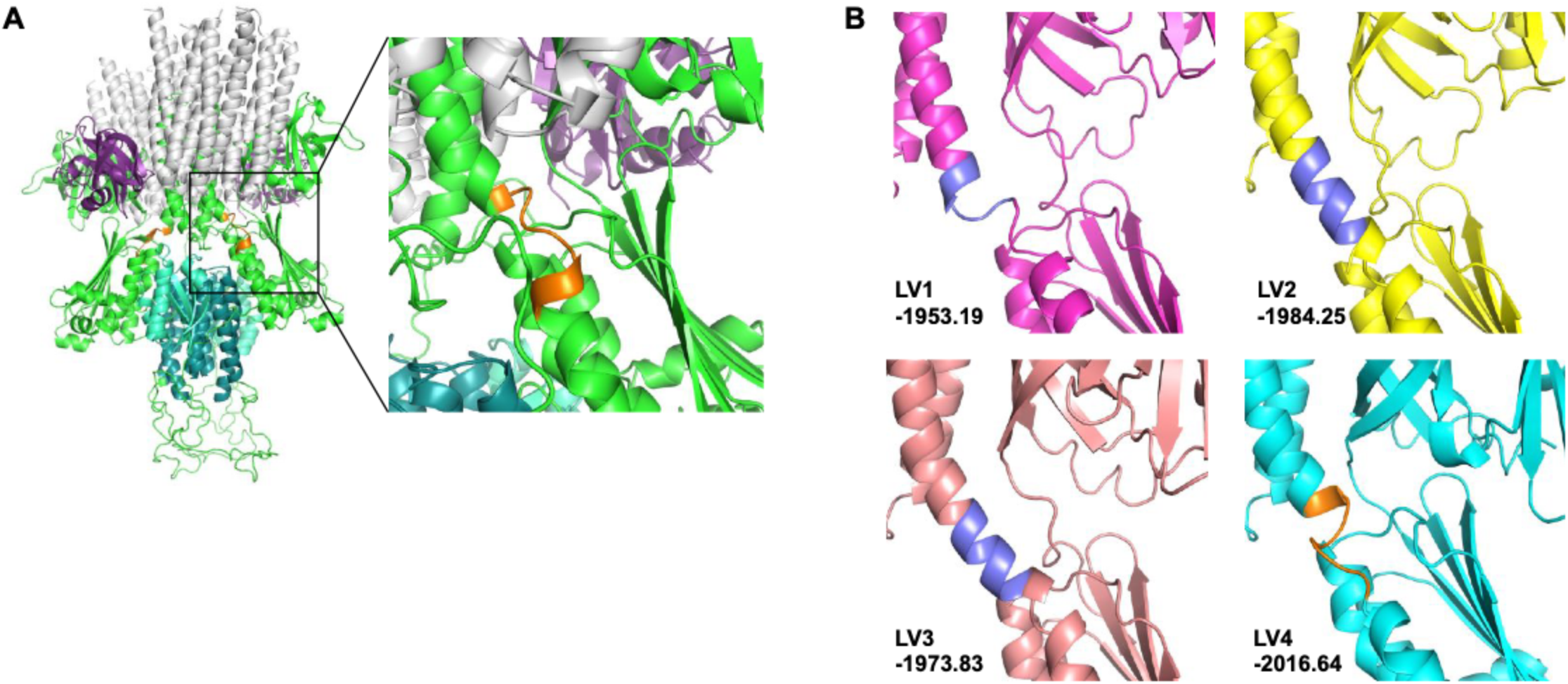
Rosetta loop modeling of L3 **A)** Model of CheA WT (P1 aquamarine, P2 cyan, P3-P5 green) in complex with CheW (purple) and Tar_F_o Short (gray) that reproduces the CheA WT inhibited conformation. The model was first reported by Muok and coworkers^31^. On the right is a close-up of L3 (orange). **(B)** The best scoring Rosetta models of CheA LV1-4 with the energies reported in REUs. α-helical structures were formed in the linker insertions AAIAAIA in LV2 and GAGAGGS in LV3. The modified linker region of LV1, GAGGS, forms an irregular loop structure. The linker insertions are colored blue for LV1-3 and the polyalanine sequence of L3 in LV4 is colored orange to match that of the WT (left). Adherence to favored Ramachandran stereochemical parameters (Rama values in REUs) of the best scoring models in the linker regions are as follows: (LV1: 0.22, LV2: -0.26, LV3:-0.13, LV4: 0.13).

### CheW Binding is Not Perturbed by Increased Linker Lengths

To test whether changes to L4 affect the folding of the P5 domain we compared binding of CheW to CheA WT and CheA-LV1. CheW (17 kDa), a paralog of P5, binds through a pseudosymmetric hydrophobic interface to the end of the P5 domain^24,36^. Chemical crosslinking with the lysine-specific reagent DSSO was employed to trap the CheA;CheW complex. CheA WT and CheA-LV1 when crosslinked run at their dimer molecular weights. Upon addition of equimolar CheW, the dimer bands shifted ∼40 kDa higher in correspondence to the crosslinked 2 CheA subunits and 2 CheW proteins **(Figure 3A)**. Thus, CheW bound to CheA-LV1 despite the extensions of the L3 and L4 linkers and it follows that the CheA P5 domain is properly folded in CheA-LV1. Although the other linker variants have the same L4 sequence as LV1, we confirmed that further extension of L3, as in Che-LV2, also did not alter CheW binding **(Figure S2)**.

**Figure 3.**
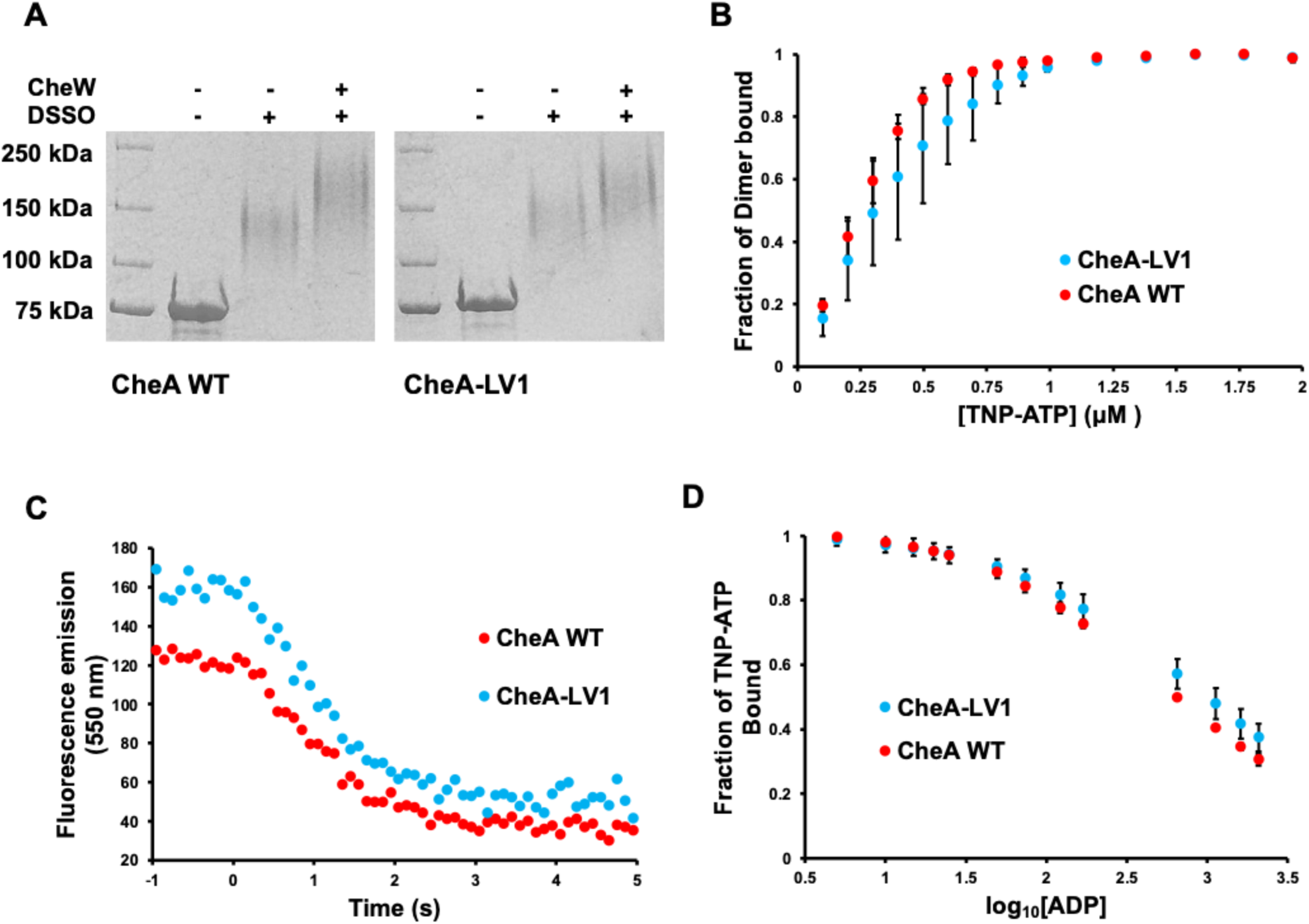
CheW and nucleotide binding by CheA LV1. **(A)** DSSO crosslinking of CheA WT and CheA-LV1 (75 kDa) to CheW (17 kDa). **(B)** TNP-ATP binding to CheA WT and CheA-LV1. Stepwise addition of TNP-ATP to CheA WT and CheA-VI gives effective IC_50_ values of 0.28 ± 0.04 and 0.4 ± 0.2 µM **(C)** ADP exchange experiment. Addition of 1000-fold molar excess of ADP to CheA WT and CheA-LV1 bound to equimolar TNP-ATP results in nucleotide exchange. Fitting of the decay curves to a first order function gave TNP-ATP k_O_ff rates of 0.45 ±0.10 and 0.40 ±0.13 1/sec for CheA WT and CheA-LV1, respectively (Number of trials; N =2). Measurement of ***_KonP∼ATP_***^rates was^ precluded by fast binding of TNP-ATP to P4. Curve shifts evident at full TNP-ATP displacement reflect instrument variation. **(D)** TNP-ATP exchange with ADP. Stepwise addition and equilibration of ADP to CheA WT and CheA-LV1 that are initially bound to TNP-ATP gives effective IC_50_ values for ADP of 400 ± 20 and 700 ± 300 µM, respectively (N = 2).

### Nucleotide Binding Affinity

To evaluate whether the reduced activity of the least active CheA-LV1 reflects a defect in ATP binding, we first compared the nucleotide binding properties of CheA-LV1 to WT with the fluorescent ATP analog TNP-ATP-(Mg^2+^). TNP-ATP-(Mg^2+^) has weak fluorescence free in solution but strong fluorescence upon binding to the CheA P4 domain^20,62^. Note that TNP-ATP is a poor phosphoryl donor to CheA^20,53,60^. TNP-ATP (and also ATP) binds to the two *Tm* CheA subunits with affinities that differ by 3-4-orders of magnitude^41^. At the concentrations of TNP-ATP and protein used, primarily one P4 domain of a CheA dimer will be occupied but some effect of the second site is also expected^20,53^. As our goal was to establish whether nucleotide binding by CheA-LV1 was substantially perturbed relative to WT, effective IC_50_ values were calculated to simplify the analysis. TNP-ATP fluorescence enhanced upon binding to CheA-LV1 and binding isotherms of CheA WT and CheA-LV1 reveal similar IC_50_ values for TNP-ATP-(Mg^2+^) **(Figure 3B)**. CheA WT and CheA-LV1 at a dimer concentration of 1 µM appear saturated at near 1 µM TNP-ATP, consistent with the tight binding expected for the first subunit and similar to that seen previously for WT^53^. To also compare dissociation rates of the nucleotides, CheA WT and CheA-LV1 were incubated with equimolar TNP-ATP, followed by an addition of 1000-fold molar excess of ADP. Loss of fluorescence as ADP was exchanged into the pocket revealed similar TNP-ATP off-rates for CheA WT and CheA-LV1 **(Figure 3C)**. In a related experiment, the concentration of TNP-ATP was held constant and ADP was titrated into solution up to a 450-fold molar excess over TNP-ATP **(Figure 3D)**. IC_50_ values calculated for ADP based on the exchange over TNP-ATP in CheA WT and CheA-LVI (405 ± 21 µM and 713 ± 308 µM, respectively) suggested only minor differences in the affinities of ADP for CheA WT and CheA-LV1 P4. Thus, extension of L3 and L4 appear to have little impact on the nucleotide affinity of P4. It follows that the P4 domain of CheA-LV1 is well folded and has a nucleotide binding pocket that resembles that of the WT kinase. That said, the slightly weaker affinities observed for both TNP-ATP and ADP by CheA-LV1 may represent a minor long-range structural effect of the linkers on nucleotide binding.

### Activity of CheA WT and Linker Variants

The autophosphorylation activities of CheA linker variants were assessed relative to CheA WT by radioisotope incorporation from γ-^32^P-ATP **(Figure 4A)**. CheA-LV1 autophosphorylation rates were only about 2% that of WT at 22 °C. The autophosphorylation activity of CheA-LV1 was similar to the level of kinase activity observed for P4 alone^30^. At 22 °C, CheA-LV2 and CheA-LV3, which extend the L3 linker further by two residues, displayed very low levels of kinase activity that were comparable to that of CheA-LV1. In contrast, CheA-LV4, which has a poly-Ala L3 sequence, gave autophosphorylation activity at 22 °C that was 23-30% of WT levels. At 55 °C, the highest tractable assay temperature for thermostable Tm CheA, activity relative to WT increased substantially for LV2, LV3 and LV4, but not LV1 **(Figure 4A)**. The fact that CheA-LV1 kinase activity at 55 °C was much less than that of CheA-LV3, which differs from CheA-LV1 only by a Gly-Ser addition, implied that the LV3 linker imparted constraints on P4, and that a helical heptad spacing favored autophosphorylation activity. CheA-LV4 activity was roughly 30% of WT at 22 °C, and all but indistinguishable from WT at 55 °C. CheA-LV4 differs from WT only in the sequence of L3, and thus despite considerable sequence conservation in L3 **(Figure 1D)** autophosphorylation activity of the isolated kinase at elevated temperatures where the Tm enzyme normally operates does not depend highly on specific L3 residues.

**Figure 4.**
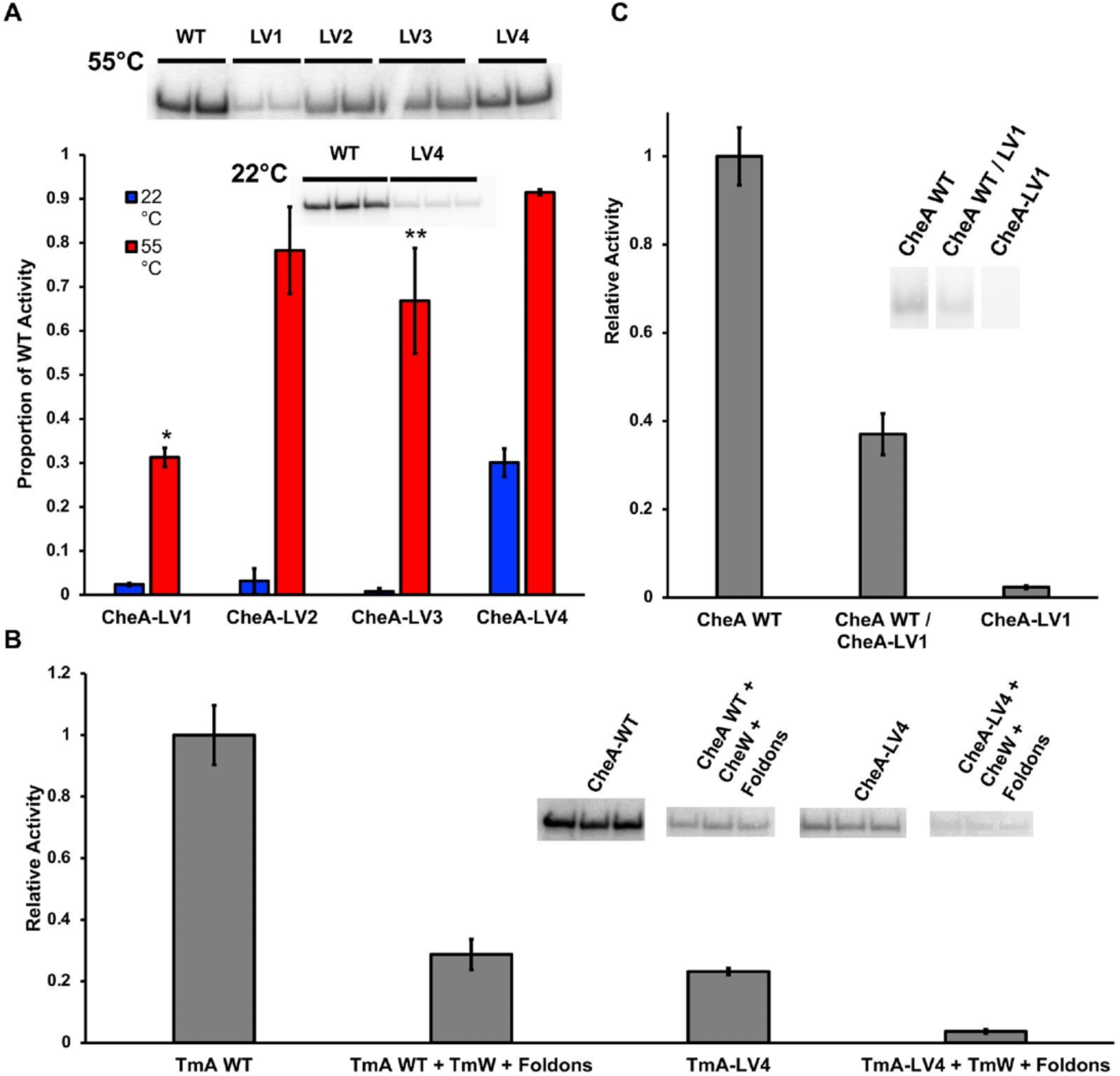
CheA autophosphorylation activity at 22 °C and 55 °C **(A)** Autophosphorylation was measured for each variant by radioisotope incorporation from γ-^32^P-ATP at 22 °C and 55 °C. Number of trials; N = 2 for all data. Error bars represent the mean deviation. *p=0.002, one-way ANOVA of CheA-LV1 compared to CheA WT. **p=0.06, one-way ANOVA of CheA-LV3 compared to CheA WT. **(B)** Autophosphorylation activity, as measured by radioisotope incorporation from γ-^32^P-ATP, of CheA WT and CheA-LV4 in the absence and presence of CheW and *E. coli* Tar_FO_ Short. The autophosphorylation assay was carried out at 22 °C. *Tm* CheA activity is reduced in the presence of receptor TOD units. CheA-LV4 also shows reduced in activity, in roughly the same proportion as WT. Insets show the relevant bands corresponding to phosphorylated CheA (N = 3). **(C)** Autophosphorylation of CheA WT, CheA-LV1, and heterodimers with mixed subunits. All samples were heated at 55°C for 10 minutes; the [P4] is the same among samples. The autophosphorylation assay was carried out at 22 °C. The CheA WT / LV1 sample contains 50% CheA WT and 50% CheA-LV1. The inset shows representative bands (N=3).

Soluble, trimeric chemoreceptor mimetics engineered from the kinase control tip region of the *E.* co//aspartate receptor Tar (Tar Foldon Short or Tar_FO_ Short) have been used to reproduce the interactions between a trimer of chemoreceptor dimers and the *Tm* CheA dimer in vitro^29,31^. As CheA-LV4 displayed appreciable kinase activity at 22 °C and only differs from CheA WT in sequence of the L3 linker, CheA-LV4 regulation by Tar Foldons may be possible. Autophosphorylation activity for free CheA-LV4 and in the presence of Tar_F0_ Short and CheW was compared to the activities of CheA WT, also free and in complex **(Figure 4B)**. Under these conditions, the Tar Foldons with CheW depressed CheA WT activity to 28% of the free kinase, whereas they depressed CheA-LV4 activity to 16%. Comparable reduction of kinase activity suggested that receptors inhibit both WT *Tm* CheA and CheA-LV4 through a similar mechanism. Owing to instability of the receptor Foldons at elevated temperatures, regulation of the other LV variants could not be tested at conditions where they are sufficiently active.

Tm CheA is active as a dimer with one P4 domain phosphorylating the P1 domain of the adjacent subunit. We therefore evaluated the effect of the LV1 linker extensions in the context of a heterodimer that contained one WT subunit and one LV1 subunit. Temperature-driven subunit exchange between *Tm* CheA dimers at 55 °C has been previously used to produce heterodimeric species^30^. *Tm* CheA WT retains its level of activity after being heated at 55 °C for 5-10 minutes and then cooled to room temperature^61^. Assuming equal exchange between equimolar WT and CheA-LV1 subunits, a statistical distribution of 1/4 CheA WT homodimer (both subunits active), 1/4 CheA-LV1 homodimer (no subunits active), and 1/2 CheA WT / CheA-LV1 heterodimer (one subunit active) would be expected **(Figure 4C)**. The roughly 37% WT activity of the heat exchanged samples (which contain 50% WT P4) indicated foremost that CheA-LV1 kinase activity is not rescued by incorporation into a CheA WT / CheA-LV1 heterodimer. Furthermore, the overall activity was slightly less than would be expected by a statistical distribution (50% activity). Thus, the CheA-LV1 subunit may reduce the activity of the WT subunit.

### Monitoring Inter-domain Interactions with Disulfide Cross-linking

Disulfide cross-linking of engineered cysteine residues has been used previously to track interdomain contacts in *Tm* CheA. For example, cross-linking between substitutions of S492C in the P4 ATP binding pocket and H45C of the substrate His residue in the P1 provide a measure of substrate domain access to the ATP-binding domain^30^. Here we similarly investigated heterodimers that contained only one P4 domain to investigate whether less restrained P4 domains within a dimer may be interfering with P1 access. Following previous protocols, subunit exchange at 55 °C between CheA S492C (WT or LV1) and CheA P1-3 H45C followed by Cu-phenanthroline-driven disulfide crosslinking and SDS-PAGE analysis of purified proteins was used to analyze for heterodimer formation^30^. CheA homodimers form inter-subunit disulfide bonds between engineered cysteine residues on their P1 or P4 domains to varying extents **(Figure 5)**. As expected, CheA FL (S492C) / P1-3 (H45C) heterodimer formation increases with heated incubation prior to cross-linking, supporting the notion that subunit exchange allows for association between S492C of CheA FL and H45C of CheA P1-3 **(Figure 5AB).** Importantly, cross-linked heterodimers formed for both LV1 and WT, thereby indicating that the inactivity of LV1 was not due to the inability of P1 to interact with a single P4 domain. Indeed, the percentage of heterodimers relative to all CheA FL species (monomers, heterodimers and homodimers) was 12 ± 8% greater for CheA LV1 compared to WT **(Figure S3)**. However, although P1 can access the P4 domain of WT and LV1, it cannot be phosphorylated in either case **(Figure 5** and **Figure S4)**. Neither a WT nor a LV1 subunit phosphorylated a P1-3 subunit in a heterodimer. Indeed, CheA autophosphorylation decreased dramatically after heat exchange with P1-3 and no P1-3 phosphorylation was observed **(Figure S4)**.

**Figure 5.**
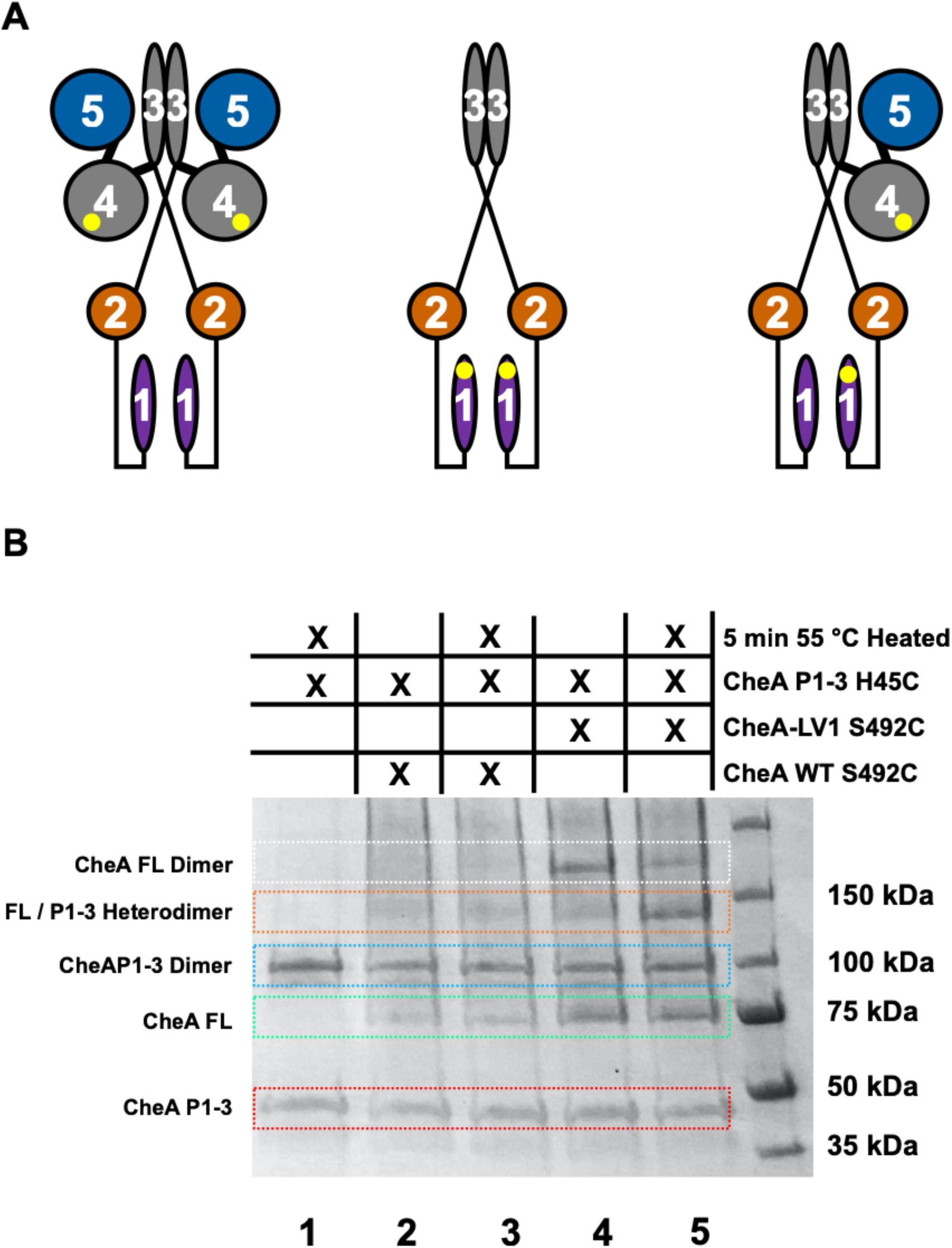
P1-to-P4 and P4-to-P4 interactions of CheA WT and CheA-LV1. **(A)** Diagrams representing CheA FL S492C homodimer, a CheA P1-3 H45C homodimer, and a CheA FL S492C / CheA P1-3 H45C heterodimer, from left to right. Yellow circles indicate cysteine substitutions. **(B)** Representative SDS-PAGE gel of disulfide crosslinking experiments. The chart above the gel specifies the contents of each sample by lane, disulfide crosslinking was initiated by the addition of Cu^2+^ (1,10)-phenanthroline. Residue 45 on P1 and 492 on P4 were changed to cysteine to report on productive substrate kinase interactions and subunit exchange was facilitated by heating samples at 55 °C for 5 min, prior to crosslinking.

### P4 Domain Proximity Correlates with Kinase Activity

The nitroxide modified nucleotide ADP-NO was employed to report on the P4-P4 intradimer distance and P4 dynamics in CheA WT and linker variants using continuous wave and pulse dipolar ESR spectroscopy (PDS) as previously described **(Figure 6A** and **Figure S5)^42^·^58^·^59^.** Cw-ESR broadening relative to free spin-label indicated binding of ADP-NO to the respective P4 domains of each CheA dimer (data not shown). Low modulation depths in the double electron electron resonance (DEER) time domain data for both CheA-LV1 and CheA WT upon incubation with ADP-NO indicated that ADP-NO did not bind tightly to both subunits, consistent with negative cooperativity in nucleotide binding between the P4 domains^41^. Nonetheless, the DEER data revealed that despite being distributed into a broad set of conformations in all cases, the ATP binding sites of the CheA-LV1 dimer were in closer proximity than in CheA WT and more closely matched the nitroxide separation in the foldon-inhibited CheA WT state **(Figure 6A)**^31^. This correlation suggests that the CheA-LV1 dimer resembles the domain arrangements of CheA when inhibited by receptors. That said, it is notable that some P4-P4 separations reported by ADP-NO in CheA-LV1 appeared even *closer* than they are in the inhibited receptor complex. These very close separations suggest association of the P4 domains, which has been observed in crystal structures of ADP-NO-bound P4 where the nitroxide moieties are flexible but could easily reside within 15 Å of each other^42^. Although these crystallographic dimers are unlikely to be represented in the full-length enzymes, alternative associations of P4 coupled with probe flexibility could produce distances in the 25-30 Å range. In contrast, CheA-LV4, the only linker variant with appreciable autophosphorylation at 22 °C, mimicked more closely the long and broad WT separation. The P4-P4 separations of LV2 and LV3 were similar to each other and intermediate to those of LV1 and LV4. LV2 and LV3 had some short components, but generally shifted to longer separations than LV1. Plotting the autophosphorylation levels of CheA WT and linker variants at 55 °C against the mean values of DEER-derived P4-P4 separation yielded a positive correlation with an R^2^ value of 0.89 **(Figure 6B)**. A similar relationship is found by plotting activity at 55 °C against the percentage of the distributions within the distance range of 0-50. Although the activities of WT, LV2, LV3 and LV4 are not greatly distinguished at 55 °C, neither are their distance distributions, unlike LV1, which has much lower activity and shorter P1-to-P1 spacing than the others.

**Figure 6.**
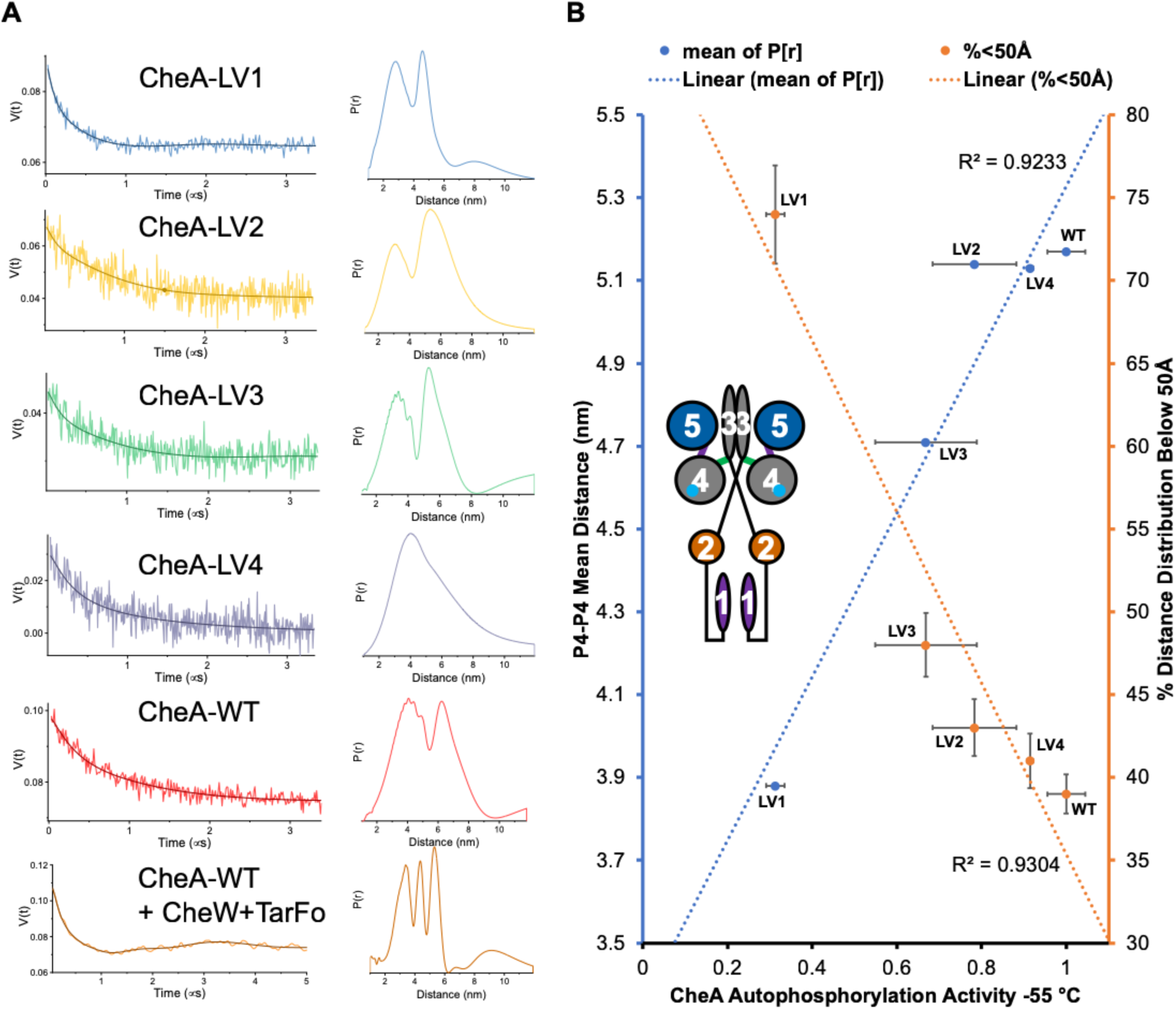
PDS of CheA variants bound to ADP-NO. **(A)** DEER-derived distance distributions of ADP-NO bound to CheA WT and linker variants. The background-corrected time-domain data of the dipolar evolution is found on the left, with reconstructed distance distributions on the left. Data for *E. coli* Tar Foldon + CheW + CheA adapted from Muok and coworkers^31^. Error analysis shown in **Figure S5**. **(B)** Plot showing correlation between autophosphorylation activity of CheA linker variants vs i) mean values of DEER-derived P4-P4 distance distributions found in (A) and ii) the percentages of each distribution than lie between 0 and 50 Â. The percentage of the CheA WT + CheW + Tar_Fo_ distance distribution, reported in (A), below 50 Å is 55 ± 6%. Distance distribution errors are derived from the distribution ranges given in **Figure S5**. Autophosphorylation activity errors represent mean deviation. Inset: a cartoon of the CheA dimer showing labeling sites (P1 ;purple, P2:brown, P34:gray, P5:blue. ADP-NO: cyan circles).

### Flexibility Analysis by SAXS

The SAXS scattering profiles of CheA WT and the linker variants indicated differences in the conformational properties of the proteins. Dimensionless Kratky plots revealed that all of the variants were substantially less compact than was the WT **(Figure 7A)**. The shift of the Kratky peaks to higher q values and shallower decays in the mid-q region for the LV proteins compared to CheA WT indicated less compact states^63-66^.

**Figure 7.**
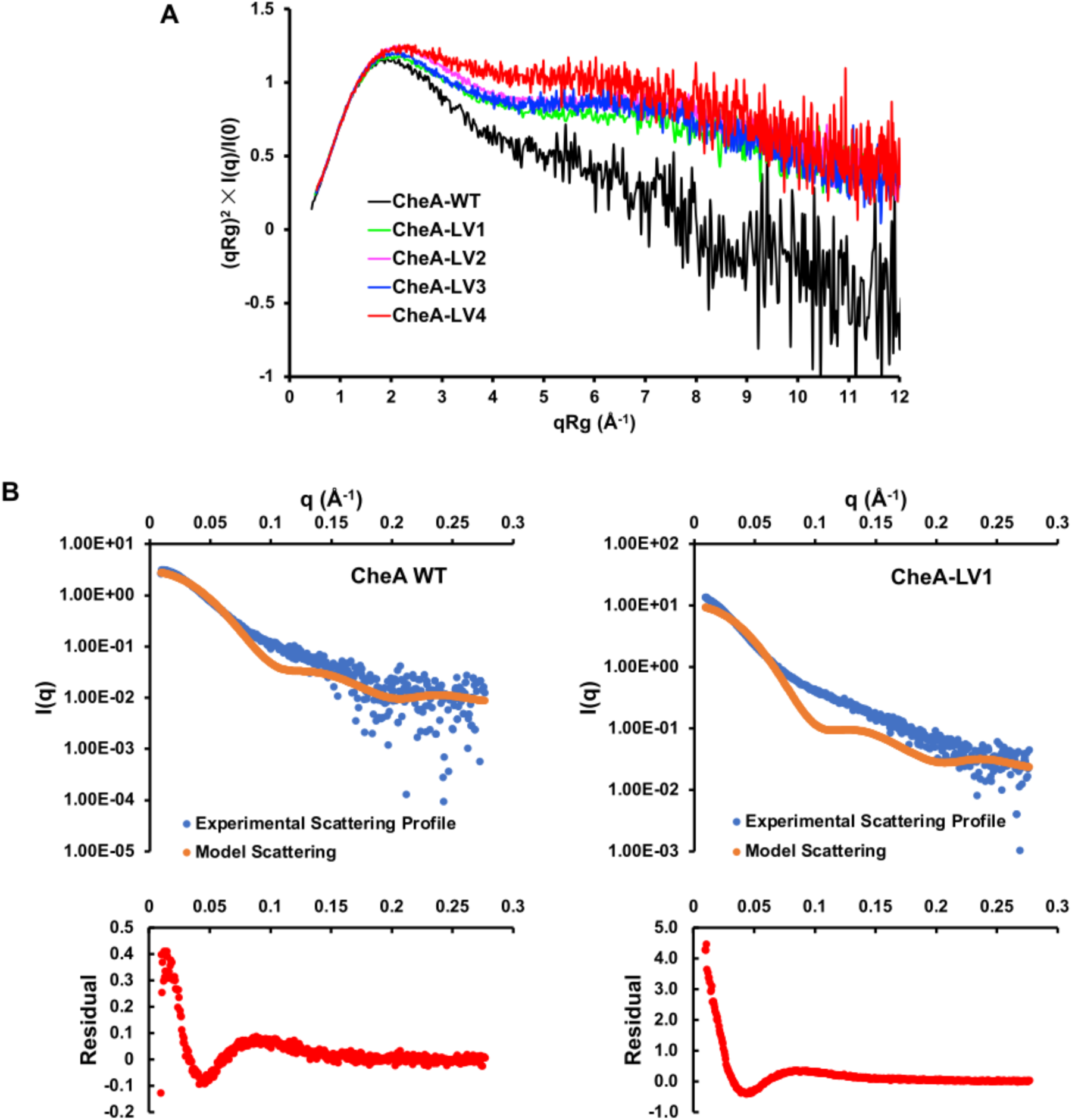
SAXS data on CheA linker variants. **(A)** Dimensionless Kratky plots of CheA WT and variants showing the mid-q region indicative of flexibility differences^30^. **(B)** Fits of experimental scattering profiles to a rigid model of inhibited CheA adapted from Muok and et al^31^. Below each fit are residuals comparing the experimentally derived and expected scattering profiles. Note the much smaller range of the residual differences displayed on the left compared to the right.

CheA-LV1, CheA-LV2, and CheA-LV3 all displayed comparable scattering profiles, of which CheA-LV1 was used as a prototype. Fitting the scattering profiles for CheA WT and CheA-LV1 to a rigid model of the kinase in its closed, inhibited state corroborated that the scattering profile of CheA WT agreed more closely with the expected scatering profile of the rigid model than did that for CheA-LV1^57^. The differences between the experimental CheA-LV1 scattering profile and the expected fit for the model suggest that increased P4 dynamics alone cannot account for the change in dynamics in CheA-LV1 **(Figure 7B)**. In particular, the PDS data discussed above showed that the P4 domains were more tightly associated in CheA-LV1 than in CheA WT and hence did not likely contribute substantially to the increased flexibility. In contrast, we have found that similar changes in protein flexibility, as monitored by SAXS, result from the increased conformational sampling of the P1, P2 and linker regions as they are released from the core kinase^30^. Thus, it appears that increases in CheA flexibility, likely associated with motion of P1, P2 and possibly L1 and L2 depend on the dynamics and interactions of the P4 domain. This finding is underscored by the behavior of CheA-LV4. CheA-LV4 had a much less compact scattering profile than CheA WT and all other linker variants, yet CheA-LV4 differs from CheA WT only in the L3 linker sequence, and in neither the L3 linker length, nor the L4 linker identity. It is quite remarkable that changes in the conformational properties of LV4 relative to WT derive from relatively modest substitutions in L3.

## Discussion

A central question in chemotaxis concerns how CheA activity is regulated by chemoreceptors. The architecture of the receptor arrays places constraints on the mechanism by which conformational changes are propagated to the CheA kinase modules, i.e. the P1 and P4 domains. The L3 and L4 linkers suspend the P4 domains below the layer composed of the receptor tips and P5:CheW rings **(Figure 1AB)** and are thus well situated to convey such conformational signals. Residue substitutions in L3 reduce both basal autophosphorylation and kinase activation by receptors, whereas changes to L4 can both decrease and increase basal autophosphorylation while reducing kinase activation^38,39^. Mechanisms for how L3 influences P4 kinase activity can be considered in two general cases. In the first case, changes to the conformations of L3 and L4 affect the properties of the catalytic components of P4^40^. In the second case, L3 and L4 control positioning and mobility of P4; linker dynamics then modulate inhibition and activation by coordinating P4-P4 interactions and by gating encounters between P4 and P1^39^. In support of case 1, single residue substitutions to L3 (M322A, R325A, M326A) decrease kinase activity^39^. Furthermore, single residue substitutions in L4 (L507A, T508A, L509A) do not support chemotaxis in vivo^39^. Additionally, changing four L4 residues in a variant devoid of the P5 domain (CheAΔPδ) can both increase and decrease kinase activity and thereby shift P4 towards active and inactive conformations^38^. In support of case 2, P1, P2 and P4 are closely associated in the inhibited form of the *Tm* kinase, activity of the P4 domains increases when dimerized by P3 and the P4 domains display negative cooperativity in binding nucleotide^30,31,53^. Our data indicate that the inhibited state of *Tm* CheA is characterized by close association of the P4 domains, but indirect effects of the linkers on the properties of the ATP binding pocket and the dynamics of the P1 substrate domain are also at play, as evidenced by the absence of CheA P1-3 phosphorylation in CheA WT / CheA P1-3 heterodimers. The P4 domains of dimeric CheA influence one another’s nucleotide binding affinity and catalytic activity as mediated by P4-P4 proximity, which is in turn controlled by L3 linker structure and dynamics.

### P4 conformation and P1-2 dynamics are coupled in FL CheA

Conformational characterization by SAXS revealed that all of the linker variants have considerably more conformational flexibility than CheA WT at room temperature, although the reduced activity of CheA-LV4 at 22 °C relative to WT demonstrates that this increase in conformational flexibility does not necessarily correlate with an increase in basal kinase activity **(Figures 6B and 7A)**. The increase in conformational dynamics of CheA-LV1 relative to CheA WT is conspicuous because the P4 domains are more tightly associated in CheA-LVI. Binding of non-hydrolyzable ATP analog 5’-adenyl(β,γ-methylene)-diphosphonate (ADPCP) to *Tm* CheA has been shown to dramatically increase CheA flexibility, the Kratky plot for which increases with positive slope and plateaus at high q-values, resembling that of a disordered Gaussian chain^30^. Such an increase in global conformational flexibility likely results from releasing regions N-terminal to P3. In a similar fashion, the Kratky plots for all of the CheA linker variants decay towards baseline but are considerably more flexible than CheA WT. Despite CheA-LV1 assuming an inhibited conformation with associated P4 domains it also shows enhanced overall domain mobility by SAXS. In a model of the inhibited CheA complex, distal regions of the P4 domains constrain the P1 domains **(Figure 2)** and thus, the decreased P4-P4 separations in LV1 may reflect the fact that the P1 domains no longer interpose the P4 domains in the inhibited state^31^. The increase in flexibility then results from greater P1, L·1, P2 or L2 conformational sampling. CheA-LV2 and CheA-LV3 have similar SAXS scattering profiles and Kratky plots as CheA-LV1. Interestingly, CheA-LV4 is even more flexible than any of the other linker variants, yet retains appreciable activity at 22 °C. Furthermore, receptor foldons inhibit CheA-LV4, thereby indicating that the dynamic nature of the full-length free protein does not prohibit inhibition by receptors. Thus, changes to L3 and L4 increased the overall flexibility of full-length CheA compared to WT; however, this increased flexibility is not reflected in greater autophosphorylation activity.

### P4-P4 proximity and CheA activity

CheA dimers are an integral feature of the receptor;kinase arrays and their component core signaling particles. In addition to the requirement of trans-subunit autophosphorylation, CheA function may depend in some manner on interactions between the two P4 domains of a dimer^30,53,67^. As the lengths and sequences of the P4 connections are perturbed in CheA-LV1 by semiflexible GSG-type linkers^68^, L3 and L4 are unlikely to be competent in propagating these signals. If the conformations of the L3 and L4 linkers play a direct role in modulating the nucleotide affinity of P4, more dramatic changes on ATP-TNP binding would likely result from the L3 and L4 variations. The proximity of the P4 ATP binding pockets in CheA-LV1, as evidenced by the PDS data and the S492C cross-linking experiments, indicated a close association of the P4 domains **(Figure 5)**. Although the PDS distributions for each variant reflected broad sampling of conformational space, the mean P4-P4 separations correlated well with autophosphorylation activity: lower activity tracks with closer P4-P4 separations. In particular, the separation distribution of the ATP-binding sites in the CheA-LV1 dimer is comparable to that of *Tm* CheA when it is inhibited by the receptor foldons. Moreover, crosslinking experiments indicate that the ATP-binding pockets face each other in some conformational states. The ability of the L3 and L4 linkers to influence P4 interactions and autophosphorylation could provide a means by which conformational signals conveyed through the linkers could regulate the kinase modules.

Although increased P4-P4 contacts correlate with reduced activity it also appears that some cooperation between the P4 domains is required for basal activity. The inability of either FL CheA-LV1 **(Figure S3A)** or FL CheA WT **(Figure S3B)** to phosphorylate CheAP1-3 despite interactions between the CheA P1-3 P1 domain and the ATP binding sites of both CheA WT and CheA-LV1 suggests that each P4 domain in a CheA dimer supports the catalytic activity of the opposing subunit. Such coupling perhaps explains why an isolated P4 domain has low activity, yet P4 activity increases substantially when it is fused to dimerized P3^30^.

### L3 linker helicity and P4 domain motion

P4 domain positioning may be a critical factor in the *Tm* CheA catalytic cycle^4,31,37^. The greater autophosphorylation of CheA-LV2-3 relative to CheA-LVI at 55 °C suggests that the P4 orientation imposed by L3 plays an important role in activity and that an extended linker can maintain this constraint if it is helical and the correct length. CheA-LV3 is longer than CheA-LV1 by only two residues, but it completes a heptad spacing that would allow a P4 domain orientation similar to that of WT if the added residues form a helix. The Rosetta modeling of the L3 elements in CheA-LV2 and CheA-LV3 indicated that those sequences do favor helical structure and that L3 in CheA-LV2 has greater helical propensity than L3 in CheA-LV3. CheA-LV4 maintains WT spacing but removes a helix-breaking Pro residue and specific side-chain contacts of the WT linker. The high activity of LV4 at elevated temperatures indicates that the WT sequence does not provide specific secondary structure or side-chain contacts that are essential for activity, however preservation of the WT L4 linker sequence in CheA-LV4 may also be responsible for the comparable activities of CheA WT and CheA-LV4. It is somewhat surprising that L3 in LV3 has the capacity to deliver structural constraints conducive to autophosphorylation given that the GSG-type sequence of the L3 extension should be quite flexible, especially at higher temperatures.

Taken together, the data indicates that the conformation of L3 controls P4-P4 proximity, which in turn influences autophosphorylation. Given that the linker variants respond differently to increases in temperature suggests that movement of the P4 domains plays a role in the catalytic cycle. Transitions between P4 conformational states has been proposed to involve a secondary structural change in L3 between a short β-strand and a kinked α-helix^31,37^. Cryo-electron tomography coupled with molecular dynamics indicate that this L3 secondary structure transition allows for critical P4 “dipping” movements but it is yet unclear how these states correlate with activity^26,37^. The catalytic defect of CheA-LV1 may result from the inability of L3 to influence P4 domain positioning and thereby prevent inhibitory P4-to-P4 interactions that become constitutive **(Figure 8)**. In contrast, the increased activity in CheA-LV2 and CheA-LV3 may derive from the ability of helical structures in these regions to separate the P4 domains. Under such a scenario, execution of the catalytic cycle includes relieving P4-P4 interactions by conformational changes in the L3 secondary structure **(Figure 8)**^68^. For LV2, LV3 and LV4, to overcome these inhibitory contacts requires increased thermal energy, available at higher temperature. Possibly the conserved Pro of the WT sequence imparts added flexibility that aids activation at low temperature. The lengthened flexible linker of CheA-LV1 releases constraints that poise the P4 domains between active and inactive states **(Figure 8)**, these constraints are then recovered in the more ordered linkers of LV2 and LV3. Our data suggests that helical structure in L3 is important for the conformational changes that relieve inhibition; however, there are several possibilities for how such a mechanism could manifest. L3 helical character in the inhibited state may be required for conversion to an active state, persist or form in a “stable” active state, or act as a transitory intermediate. An added complication is that the two P4 domains appear to work together to achieve an active state, as suggested by the P1-3 heterodimer phosphotransfer experiments. This cooperativity may involve direct interactions of the two P4 domains that differ from those of the inhibited state, but could also arise from their coordinated movement, which may be essential for binding and release of one or both of ATP and P1^21^·^69^. The active state of CheA may not only involve P4 dynamics, but also P4 asymmetry. Surprisingly, changes in P4 conformation also affect the P1 and P2 domains, which become freely mobile in all the linker variants, presumably because the domain interactions found in the WT are perturbed. Although CheA has long been considered a protein composed of domains with specific dedicated functions^4^, the dimeric enzyme appears to function as a cooperative unit, with the L3 and L4 linkers playing critical roles in the cooperative interactions required for autophosphorylation.

**Figure 8.**
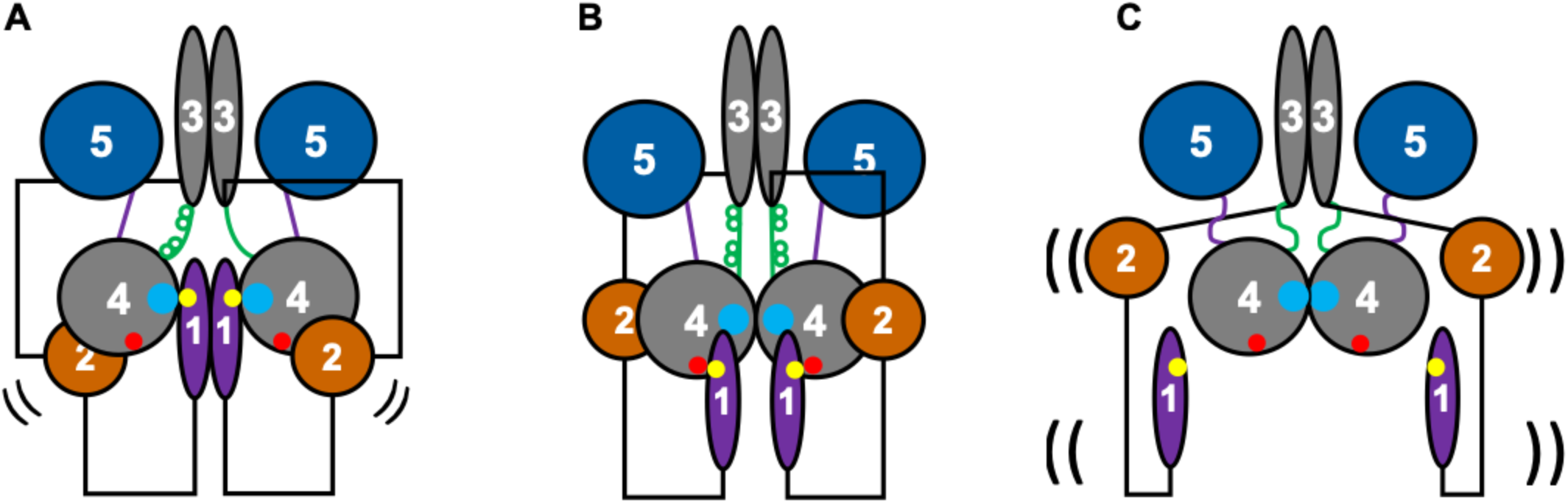
Effects of the P4 linkers on CheA Structure and Dynamics. **(A)** In the active form of CheA WT, the L3 linker structure (green), an α-helix interrupted by a proline residue in the WT sequence **(Fig. 1D)**, facilitates separation of the P4 domains from one another to allow access of the P1 substrate His residue (yellow dot) to the P4 ATP binding site (blue dot). P4 domain motions important for autophosphorylation may require L3 helical structure but could also involve conformational change between an α-helix and a β-strand^37^, as indicated in one subunit (See Discussion). The active state could be symmetric or asymmetric. The associations among P1, P2, and P4 are not as tight as those found in the receptor inhibited complex and involve a more productive association between P1 and P4 than in the receptor-inhibited state in (B). Hence, the active state may be termed as “open” only in the sense that the domain motions become more dynamic and their interactions more transient. Flexibility relative to (B) and (C) is indicated by black arcs. **(B)** The receptor-inhibited CheA WT state characterized by an inactive P4-P4 dimer that sequesters the P4 ATP binding pockets. P1 binds to a P4 inhibitory site (red dot)^4,70^. **(C)** The inhibited state of CheA-LV1. Flexibility relative to (A) and (B) is indicated by black arcs. The L3 and L4 linkers are extended and flexible. Failure to modulate P4-P4 separation due to the extended L3 linker results in an inactive P4 dimer. Occlusion of P1 from P4 results in P1-2 release and greater overall dynamics, but no productive interactions between P1 and P4.

#### Supporting Information Available

SEC traces of CheA and linker variants; DSSO crosslinking of LV2; SDS-PAGE gels of heterodimer crosslinking experiments and analysis; autophosphorylation experiments; PDS distance distributions with error bounds.

## Acknowledgements

We thank Dr. Alise R. Muok for valuable discussions, Joanne Widom for plasmid construction, Yajie Xu for help with CheW binding experiments, and the Cornell High Energy Synchrotron Source (CHESS) and the National Biomedical Center for Advanced ESR Technologies (ACERT) for access to data collection facilities. This work was supported by grants from the National Institutes of Health: R35GM122535 and R35GMR024 to BRC. ACERT is supported by P41GM103521. CHESS is supported by NSF award DMR-1332208 and NIH/NIGMS award P30GM103485.

**Accession IDs of Proteins from UniProt**

*Thermotoga maritima* CheY: Q56312

*Thermotoga maritima* CheW: Q56311

*Thermotoga maritima* CheA

## Supporting Information

**Supplemental Figure 1.**
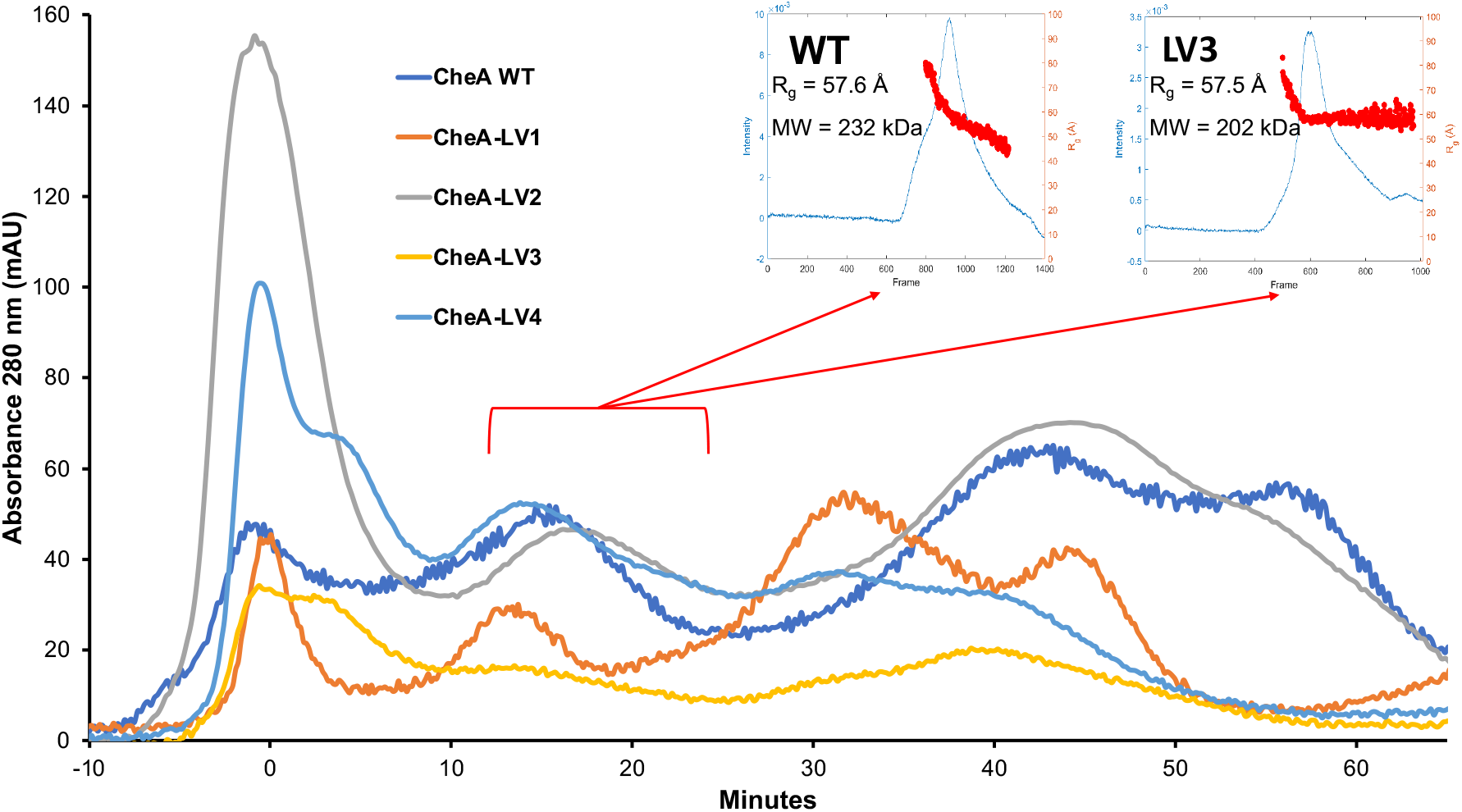
Size-exclusion chromatography traces of CheA WT and linker variants. Each of the proteins was purified on a Preparative Superdex 200 column. The void peaks were aligned at 0 min to account for variations in column performance. The red bracket indicates the SEC peak collected for the dimer species. These proteins were then used in SEC-SAXS, representative data for which is displayed in insets. MW estimations from the Porod volume, as well as radius of gyration (Rg) measuerments indicate that the CheA WT and linker variants are dimeric.

**Supplementary Figure 2.**
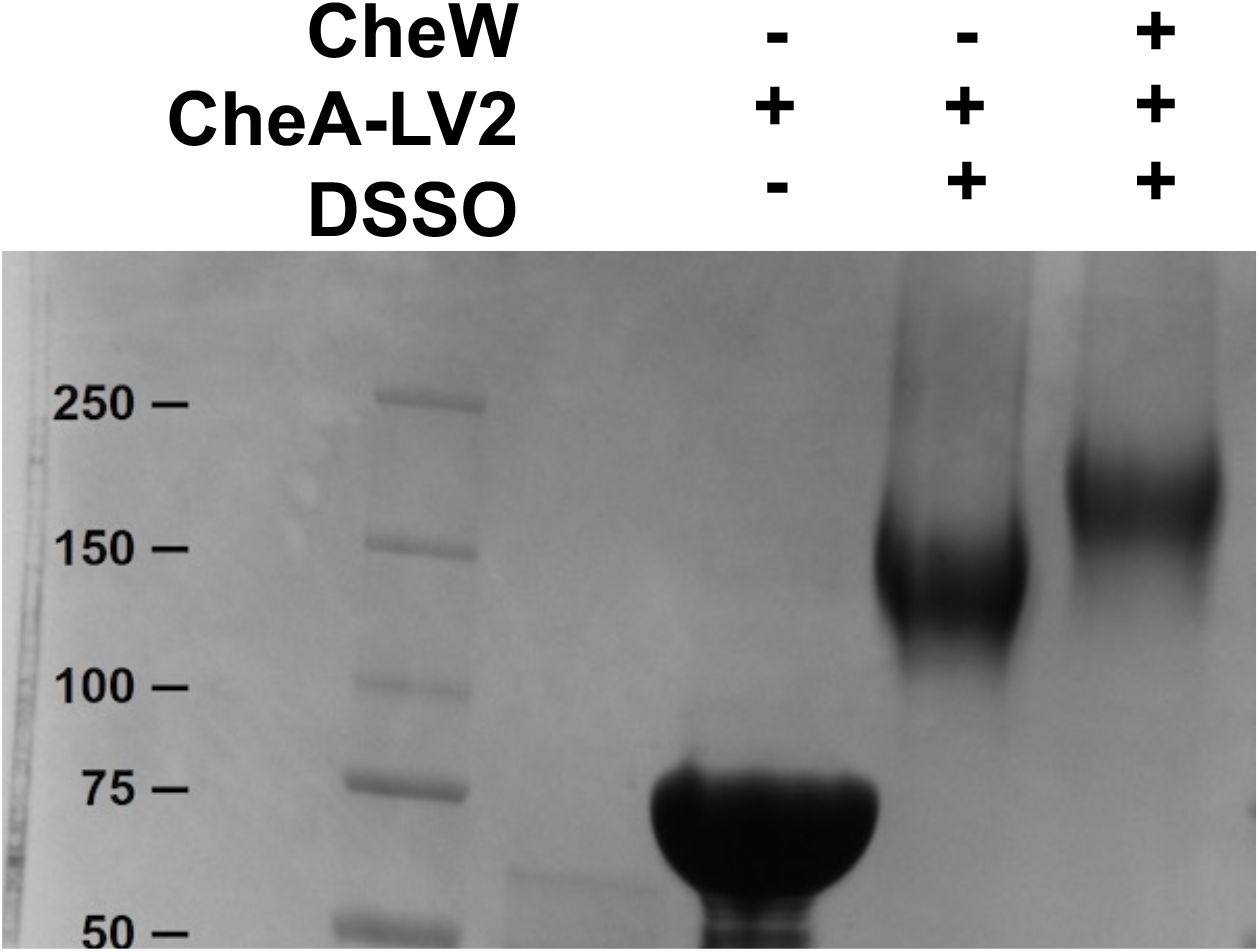
DSSO crosslinking of CheA-LV2 (75 kDa) to CheW (17 kDa). The molecular weight markers are indicated in kilodaltons (kDa) to the left.

**Supplemental Figure 3.**
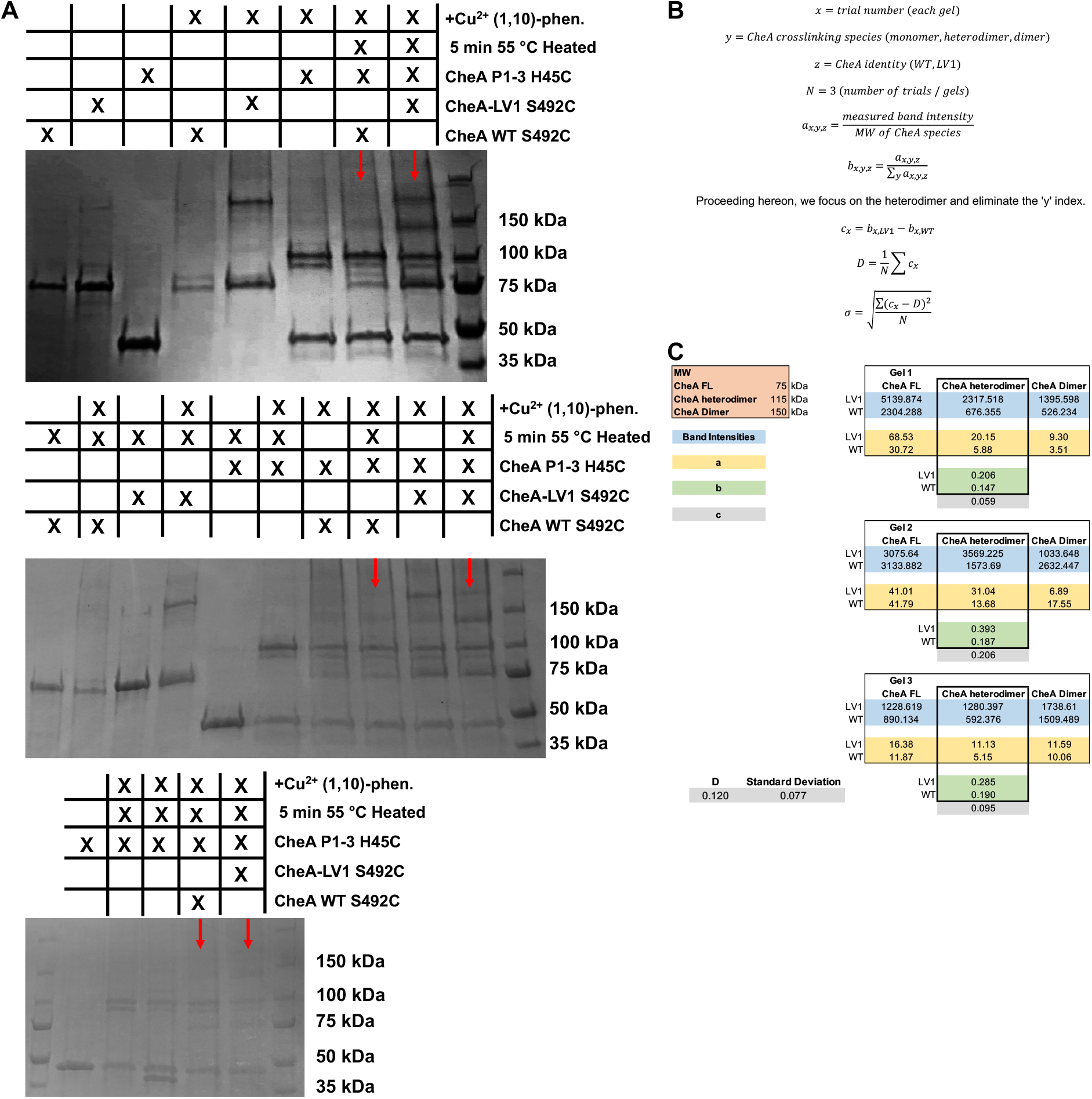
Quantification of CheA FL / CheA P1-3 Heterodimer Formation **(A)** SDS-PAGE gels of disulfide crosslinking experiments used to quantify CheA FL bands. Red arrows indicate the lanes used in that calculation. The chart above each gel specifies the contents of each sample by lane, cysteine-cysteine crosslinking was initiated by the addition of Cu^2+^ (1,10)-phenanthroline. Subunit exchange was provided by heating samples at 55 °C for 5 mins. **(B)** Detail of calculations shown in (C). First, the band intensities of crosslinking products measured in ImageJ are corrected "a" for their molecular weights. Next, the proportions "b" of each CheA FL crosslinking species are calculated. The percent increases "c" of the heterodimer band for LV1 relative to WT is calculated, and their average "D" is reported with its standard deviation. **(C)** Raw data and outcome of calculations detailed in (B). The summary shown at the bottom left reveals that the percentage of CheA FL / CheA P1-3 heterodimer in CheA-LV1 increases 12 ± 8 % relative to CheA WT.

**Supplemental Figure 4.**
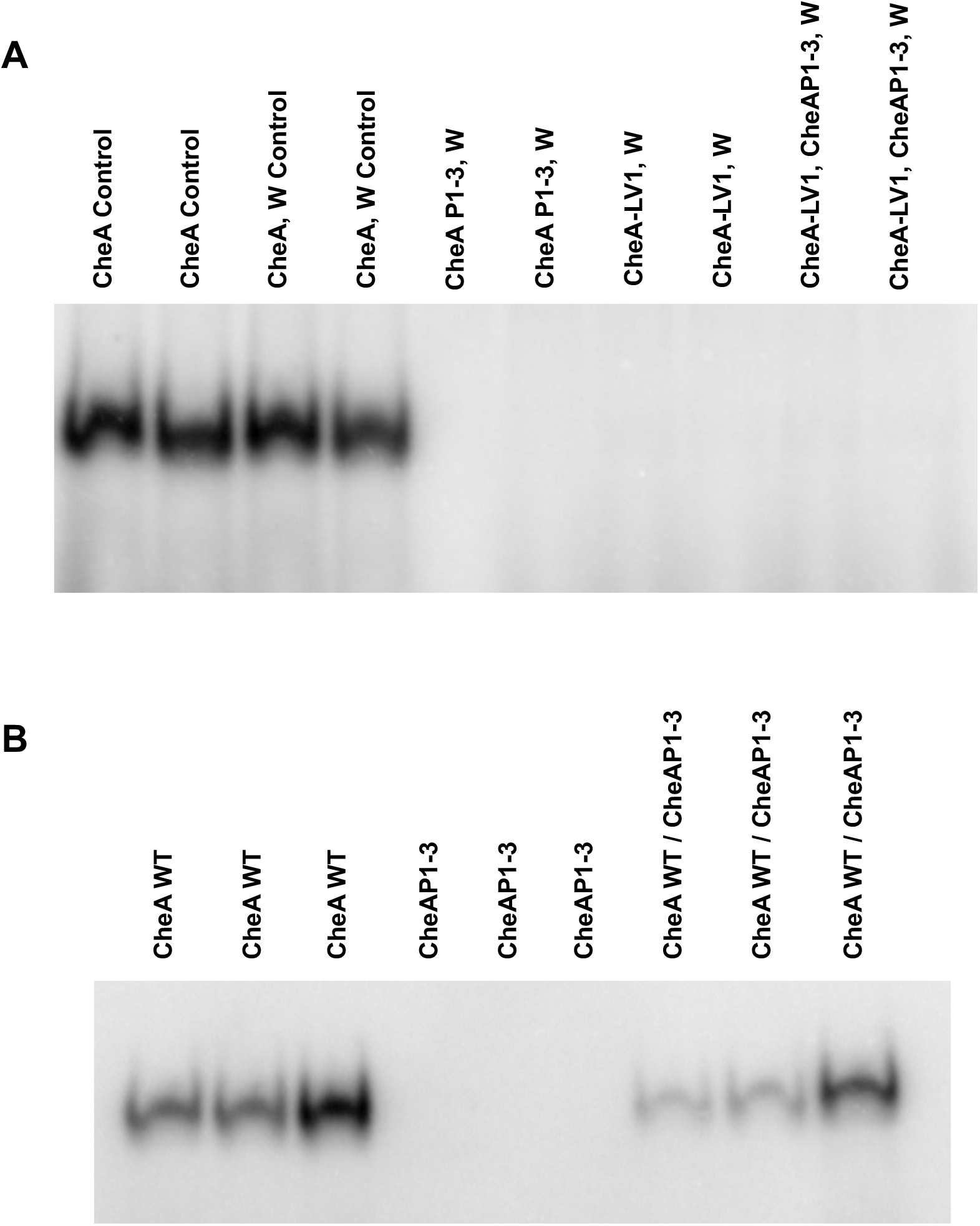
Autophosphorylation of CheA WT and CheA-LV1 after CheAP1-3 subunit exchange. **(A)** Autoradiography of WT and CheA-LV1 autophosphorylation with and without CheAP1-3 after subunit exchange. CheA-LV1 registers at very low autophosphorylation. CheAP1-3 bands are absent. **(B)** Autoradiography of CheA WT autophosphorylation with and without CheAP1-3 after subunit exchange. Phosphorylated CheAP1-3 bands are absent in all cases and CheAP1-3 reduces WT CheA autophosphorylation after subunit exchange.

**Supplemental Figure 5.**
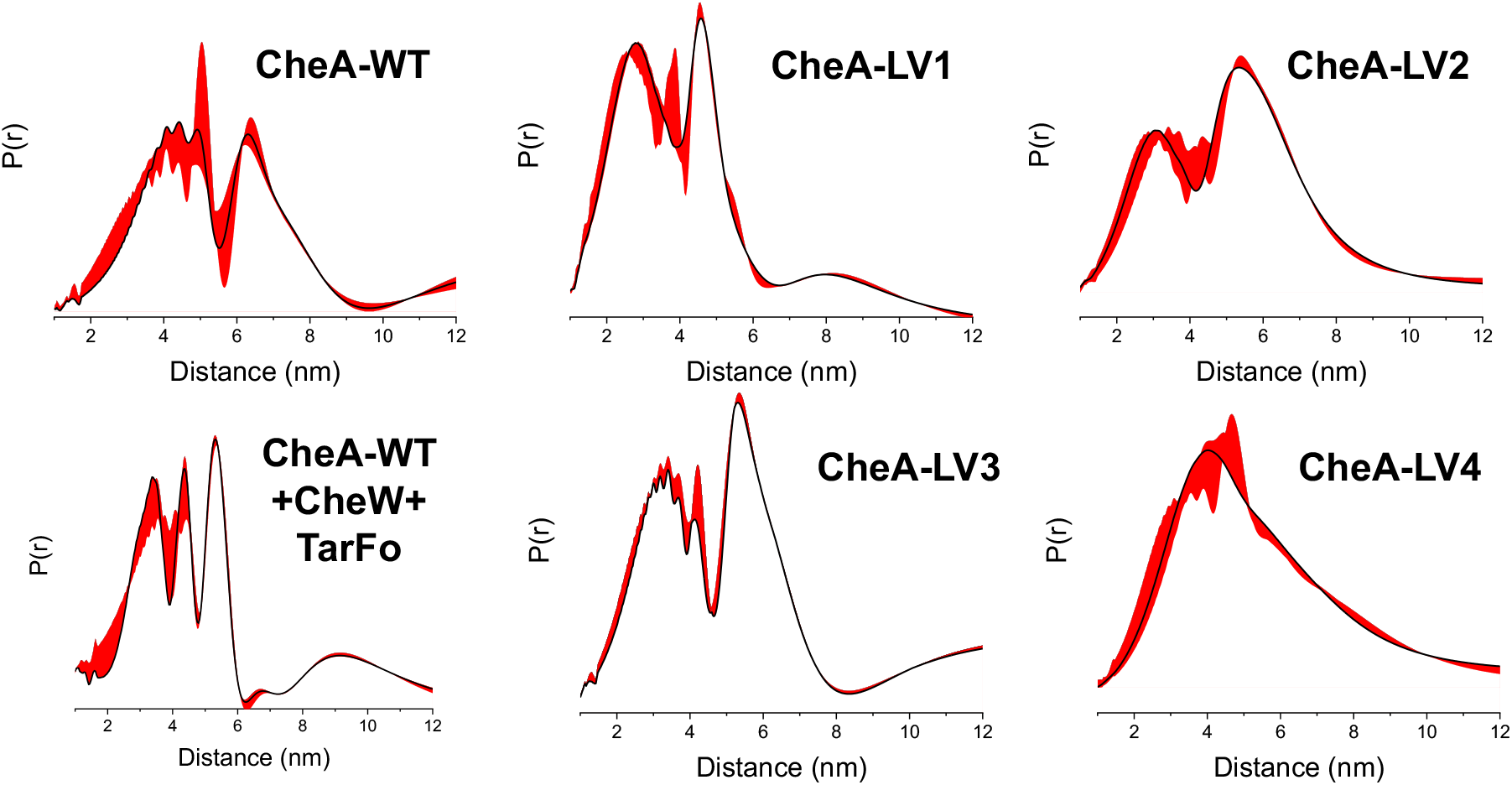
Distance distribution reconstructions by WavPDS with error bounds (red) for ADP-NO bound WT CheA, CheA WT bound to CheW and Tar**Fo** and CheA linker variants^58^.

## References

(1) Armitage, J. P. Bacterial Tactic Responses. In Advances in Microbial Physiology, Elsevier, 1999; Vol. 41, pp 229–289. https://doi.org/10.1016/S0065-2911(08)60168-X

(2) Parkinson, J. S.; Hazelbauer, G. L.; Falke, J. J. Signaling and Sensory Adaptation in Escherichia Coli Chemoreceptors: 2015 Update. Trends in Microbiology 2015, 23 (5), 257–266. https://doi.Org/10.1016/j.tim.2015.03.003.

(3) Hazelbauer, G. L.; Falke, J. J.; Parkinson, J. S. Bacterial Chemoreceptors: High-Performance Signaling in Networked Arrays. Trends in Biochemical Sciences 2008, 33 (1), 9–19. https://doi.Org/10.1016/j.tibs.2007.09.014.

(4) Muok, A. R.; Briegel, A.; Crane, B. R. Regulation of the Chemotaxis Histidine Kinase CheA: A Structural Perspective. Biochimica et Biophysica Acta (BBA) - Biomembranes 2020, 1862 (1), 183030. https://doi.Org/10.1016/j.bbamem.2019.183030.

(5) Sultan, S. Z.; Manne, A.; Stewart, P. E.; Bestor, A.; Rosa, P. A.; Charon, N. W.; Motaleb, M. A. Motility Is Crucial for the Infectious Life Cycle of Borrelia Burgdorferi. Infect Immun 2013, 81 (6), 2012–2021. https://doi.org/10.1128/IAI.01228-12.

(6) Rolig, A. S.; Carter, J. E.; Ottemann, K. M. Bacterial Chemotaxis Modulates Host Cell Apoptosis to Establish a T-Helper Cell, Type 17 (Th17)-Dominant Immune Response in Helicobacter Pylori Infection. Proceedings of the National Academy of Sciences 2011, 708(49), 19749-19754. https://doi.org/10.1073/pnas.1104598108.

(7) Howitt, M. R.; Lee, J. Y.; Lertsethtakarn, P.; Vogelmann, R.; Joubert, L.-M.; Ottemann, K. M.; Amieva, M. R. ChePep Controls Helicobacter Pylori Infection of the Gastric Glands and Chemotaxis in the *Epsilonproteobacteria*. mBio 2011, 2 (4). https://doi.org/10.1128/mBio.00098-11.

(8) Kroupitski, Y.; Golberg, D.; Belausov, E.; Pinto, R.; Swartzberg, D.; Granot, D.; Sela, S. Internalization of Salmonella Enterica in Leaves Is Induced by Light and Involves Chemotaxis and Penetration through Open Stomata. AEM 2009, 75 (19), 6076–6086. https://doi.org/10.1128/AEM.01084-09.

(9) Sze, C. W.; Zhang, K.; Kariu, T.; Pal, U.; Li, C. Borrelia Burgdorferi Needs Chemotaxis To Establish Infection in Mammals and To Accomplish Its Enzootic Cycle. Infect. Immun. 2012, 80 (7), 2485–2492. https://doi.org/10.1128/IAI.00145-12.

(10) Croxen, M. A.; Sisson, G.; Melano, R.; Hoffman, P. S. The Helicobacter Pylori Chemotaxis Receptor TIpB (HP0103) Is Required for PH Taxis and for Colonization of the Gastric Mucosa. Jβ 2006, 788(7), 2656–2665. https://doi.Org/10.1128/JB.188.7.2656-2665.2006.

(11) Hazelbauer, G. L.; Lai, W. C. Bacterial Chemoreceptors: Providing Enhanced Features to Two-Component Signaling. Current Opinion in Microbiology 2010, 13 (2), 124–132. https://doi.Org/10.1016/j.mib.2009.12.014.

(12) Wadhams, G. H.; Armitage, J. P. Making Sense of It All: Bacterial Chemotaxis. Nat.Rev.Mol.Cell Biol. 2004, 5(12), 1024–1037. https://doi.org/10.1038/nrm1524.

(13) Bilwes, A. M.; Alex, L. A.; Crane, B. R.; Simon, M. I. Structure of CheA, a Signal-Transducing Histidine Kinase. 11.

(14) Gloor, S. L.; Falke, J. J. Thermal Domain Motions of CheA Kinase in Solution: Disulfide Trapping Reveals the Motional Constraints Leading to Trans-Autophosphorylation. 14.

(15) Morrison, T. B.; Parkinson, J. S. Liberation of an Interaction Domain from the Phosphotransfer Region of CheA, a Signaling Kinase of Escherichia Coli. Proceedings of the National Academy of Sciences 1994, 91 (12), 5485–5489. https://doi.org/10.1073/pnas.91.12.5485.

(16) Stewart, R. C.; Jahreis, K.; Parkinson, J. S. Rapid Phosphotransfer to CheY from a CheA Protein Lacking the CheY-Binding Domain t. Biochemistry 2000, 39 (43), 13157–13165. https://doi.org/10.1021/bi001100k.

(17) Borkovich, K. A.; Kaplan, N.; Hess, J. F.; Simon, M. I. Transmembrane Signal Transduction in Bacterial Chemotaxis Involves Ligand-Dependent Activation of Phosphate Group Transfer. Proceedings of the National Academy of Sciences 1989, 86 (4), 1208–1212. https://doi.Org/10.1073/pnas.86.4.1208.

(18) Bourret, R. B.; Borkovich, K. A.; Simon, M. I. Signal Transduction Pathways Involving Protein Phosphorylation in Prokaryotes. Annu. Rev. Biochem. 1991, 60 (1), 401–441. https://doi.org/l0.1146/annurev.bi.60.070191.002153.

(19) Levit, M. N.; Liu, Y.; Stock, J. B. Mechanism of CheA Protein Kinase Activation in Receptor Signaling Complexes. Biochemistry 1999, 38 (20), 6651–6658. https://doi.org/10.1021/bi982839l.

(20) Eaton, A. K.; Stewart, R. C. Kinetics of ATP and TNP-ATP Binding to the Active Site of CheA from *Thermotoga Maritima*. Biochemistry 2010, 49 (27), 5799–5809. https://doi.org/10.1021/bi100721b.

(21) Jun, S.-Y.; Pan, W.; Hazelbauer, G. L. ATP Binding as a Key Target for Control of the Chemotaxis Kinase. Journal of Bacteriology 2020, 202 (13), 13.

(22) Mello, B. A.; Pan, W.; Hazelbauer, G. L.; Tu, Y. A Dual Regulation Mechanism of Histidine Kinase CheA Identified by Combining Network-Dynamics Modeling and System-Level Input-Output Data. PLoS Comput Biol 2018, 14 (7), e1006305. https://doi.org/10.1371/journal.pcbi.1006305.

(23) Hess, J. F.; Oosawa, K.; Kaplan, N.; Simon, M. I. Phosphorylation of Three Proteins in the Signaling Pathway of Bacterial Chemotaxis. Cell 1988, 53 (1), 79–87. https://doi.org/l0.1016/0092-8674(88)90489-8.

(24) Briegel, A.; Li, X.; Bilwes, A. M.; Hughes, K. T.; Jensen, G. J.; Crane, B. R. Bacterial Chemoreceptor Arrays Are Hexagonally Packed Trimers of Receptor Dimers Networked by Rings of Kinase and Coupling Proteins. Proceedings of the National Academy of Sciences of the United States of America 2012, 109 (10), 3766–3771. https://doi.Org/10.1073/pnas.1115719109.

(25) Liu, J.; Hu, B.; Morado, D. R.; Jani, S.; Manson, M. D.; Margolin, W. Molecular Architecture of Chemoreceptor Arrays Revealed by Cryoelectron Tomography of Escherichia Coli Minicells. Proceedings of the National Academy of Sciences 2012, 109 (23), E1481–E1488. https://doi.org/10.1073/pnas.1200781109.

(26) Cassidy, C. K.; Himes, B. A.; Alvarez, F. J.; Ma, J.; Zhao, G.; Perilla, J. R.; Schulten, K.; Zhang, P. CryoEM and Computer Simulations Reveal a Novel Kinase Conformational Switch in Bacterial Chemotaxis Signaling. eLife 2015, 4, e08419. https://doi.org/10.7554/eLife.08419.

(27) Piñas, G. E.; DeSantis, M. D.; Cassidy, C. K.; Parkinson, J. S. Hexameric Rings of the Scaffolding Protein CheW Enhance Response Sensitivity and Cooperativity in *Escherichia Coli* Chemoreceptor Arrays. Sc/. Signal. 2022, 15 (718), eabj1737. https://doi.org/l0.1126/scisignal.abj1737.

(28) Borkovich, K. A.; Simon, M. I. The Dynamics of Protein Phosphorylation in Bacterial Chemotaxis. Ce//1990, 63 (6), 1339–1348. https://doi.org/10.1016/0092-8674(90)90429-1.

(29) Greenswag, A. R.; Li, X.; Borbat, P. P.; Samanta, D.; Watts, K. J.; Freed, J. H.; Crane, B. R. Preformed Soluble Chemoreceptor Trimers That Mimic Cellular Assembly States and Activate CheA Autophosphorylation. Biochemistry 2015, 54 (22), 3454–3468. https://doi.org/10.1021/bi501570n.

(30) Greenswag, A. R.; Muok, A.; Li, X.; Crane, B. R. Conformational Transitions That Enable Histidine Kinase Autophosphorylation and Receptor Array Integration. Journal of Molecular **β/o/ogy** 2015, 427(24), 3890–3907. https://doi.Org/10.1016/j.jmb.2015.10.015.

(31) Muok, A. R.; Chua, T. K.; Srivastava, M.; Yang, W.; Maschmann, Z.; Borbat, P. P.; Chong, J.; Zhang, S.; Freed, J. H.; Briegel, A.; Crane, B. R. Engineered Chemotaxis Core Signaling Units Indicate a Constrained Kinase-off State. Sci. Signal. 2020, 13 (657), eabc1328. https://doi.org/l0.1126/scisignal.abc1328.

(32) Li, X.; Fleetwood, A. D.; Bayas, C.; Bilwes, A. M.; Ortega, D. R.; Falke, J. J.; Zhulin, I. B.; Crane, B. R. The 3.2 Å Resolution Structure of a Receptor;CheAΌheW Signaling Complex Defines Overlapping Binding Sites and Key Residue Interactions within Bacterial Chemosensory Arrays. 2013, 14.

(33) Merz, G. E.; Borbat, P. P.; Muok, A. R.; Srivastava, M.; Bunck, D. N.; Freed, J. H.; Crane, B. R. Site-Specific Incorporation of a Cu ^2+^ Spin Label into Proteins for Measuring Distances by Pulsed Dipolar Electron Spin Resonance Spectroscopy. J. Phys. Chem. B 2018, 122 (41), 9443–9451. https://doi.org/10.1021/acs.jpcb.8b05619.

(34) Briegel, A.; Ames, P.; Gumbart, J. C.; Oikonomou, C. M.; Parkinson, J. S.; Jensen, G. J. The Mobility of Two Kinase Domains in the *E Scherichia Coli* Chemoreceptor Array Varies with Signalling State. Molecular Microbiology 2013, 89 (5), 831–841. https://doi.Org/10.1111/mmi.12309.

(35) Pan, W.; Dahlquist, F. W.; Hazelbauer, G. L. Signaling Complexes Control the Chemotaxis Kinase by Altering Its Apparent Rate Constant of Autophosphorylation: Control of the Chemotaxis Kinase. Protein Science 2017, 26 (8), 1535–1546. https://doi.Org/10.1002/pro.3179.

(36) Piñas, G. E.; DeSantis, M. D.; Parkinson, J. S. Noncritical Signaling Role of a Kinase-Receptor Interaction Surface in the Escherichia Coli Chemosensory Core Complex. Journal of Molecular β/o/ogy 2018, 430(7), 1051–1064. https://doi.org/10.1016/jjmb.2018.02.004.

(37) Cassidy, C. K.; Himes, B. A.; Sun, D.; Ma, J.; Zhao, G.; Parkinson, J. S.; Stansfeld, P. J.; Luthey-Schulten, Z.; Zhang, P. Structure and Dynamics of the E. Coli Chemotaxis Core Signaling Complex by Cryo-Electron Tomography and Molecular Simulations. Commun Biol 2020, 3(1), 24. https://doi.org/10.1038/s42003-019-0748-0.

(38) Ding, X.; He, Q.; Shen, F.; Dahlquist, F. W.; Wang, X. Regulatory Role of an Interdomain Linker in the Bacterial Chemotaxis Histidine Kinase CheA. J Bacteriol 2018, 200 (10), e00052–18. https://doi.org/10.1128/JB.00052-18.

(39) Wang, X.; Wu, C.; Vu, A.; Shea, J.-E.; Dahlquist, F. W. Computational and ExperimentalAnalyses Reveal the Essential Roles of Interdomain Linkers in the Biological Function of Chemotaxis Histidine Kinase CheA. J. Am. Chem. Soc. 2012, 134 (39), 16107–16110. https://doi.org/10.1021/ja3056694.

(40) Wang, X.; Vallurupalli, P.; Vu, A.; Lee, K.; Sun, S.; Bai, W.-J.; Wu, C.; Zhou, H.; Shea, J.-E.; Kay, L. E.; Dahlquist, F. W. The Linker between the Dimerization and Catalytic Domains of the CheA Histidine Kinase Propagates Changes in Structure and Dynamics That Are Important for Enzymatic Activity. Biochemistry 2014, 53 (5), 855–861. https://doi.org/10.1021/bi4012379.

(41) Eaton, A. K.; Stewart, R. C. The Two Active Sites of *Thermotoga Maritima* CheA Dimers Bind ATP with Dramatically Different Affinities. Biochemistry 2009, 48 (27), 6412–6422. https://doi.org/10.1021/bi900474g.

(42) Muok, A. R.; Chua, T. K.; Le, H.; Crane, B. R. Nucleotide Spin Labeling for ESR Spectroscopy of ATP-Binding Proteins. Appl Magn Reson 2018, 49 (12), 1385–1395. https://doi.org/l0.1007/S00723-018-1070-6.

(43) Mandell, D. J.; Coutsias, E. A.; Kortemme, T. Sub-Angstrom Accuracy in Protein Loop Reconstruction by Robotics-Inspired Conformational Sampling. Nat Methods 2009, 6 (8), 551–552. https://doi.org/l0.1038/nmeth0809-551.

(44) Wang, C.; Bradley, P.; Baker, D. Protein-Protein Docking with Backbone Flexibility. Journal of Molecular Biology 2007, 373 (2), 503–519. https://doi.Org/10.1016/j.jmb.2007.07.050.

(45) Conway, P.; Tyka, M. D.; DiMaio, F.; Konerding, D. E.; Baker, D. Relaxation of Backbone Bond Geometry Improves Protein Energy Landscape Modeling: Relaxation of Backbone Bond Geometry. Protein Science 2014, 23 (1), 47–55. https://doi.org/10.1002/pro.2389.

(46) Khatib, F.; Cooper, S.; Tyka, M. D.; Xu, K.; Makedon, I.; Popovic, Z.; Baker, D.; Players, F. Algorithm Discovery by Protein Folding Game Players. Proceedings of the National Academy of Sciences 2011, 708(47), 18949–18953. https://doi.org/10.1073/pnas.1115898108.

(47) Tyka, M. D.; Keedy, D. A.; André, I.; DiMaio, F.; Song, Y.; Richardson, D. C.; Richardson, J. S.; Baker, D. Alternate States of Proteins Revealed by Detailed Energy Landscape Mapping. Journal of Molecular Biology 2011, 405 (2), 607–618. https://doi.Org/10.1016/j.jmb.2010.11.008.

(48) Hirst, S. J.; Alexander, N.; Mchaourab, H. S.; Meiler, J. RosettaEPR: An Integrated Tool for Protein Structure Determination from Sparse EPR Data. Journal of Structural Biology 2011, 173 (3), 506–514. https://doi.Org/10.1016/j.jsb.2010.10.013.

(49) Alexander, N.; Al-Mestarihi, A.; Bortolus, M.; Mchaourab, H.; Meiler, J. De Novo High-Resolution Protein Structure Determination from Sparse Spin-Labeling EPR Data. Structure 2008, 76(2), 181–195. https://doi.Org/10.1016/j.str.2007.11.015.

(50) Huang, P.-S.; Ban, Y.-E. A.; Richter, F.; Andre, I.; Vernon, R.; Schief, W. R.; Baker, D. RosettaRemodel: A Generalized Framework for Flexible Backbone Protein Design. PLoS ONE 2011, 6(8), e24109. https://doi.org/10.1371/journal.pone.0024109.

(51) Hughson, A. G.; Hazelbauer, G. L. Detecting the Conformational Change of Transmembrane Signaling in a Bacterial Chemoreceptor by Measuring Effects on Disulfide Cross-Linking in Vivo. Proceedings of the National Academy of Sciences 1996, 93 (21), 11546–11551. https://doi.org/10.1073/pnas.93.21.11546.

(52) Kao, A.; Chiu, C.; Vellucci, D.; Yang, Y.; Patel, V. R.; Guan, S.; Randall, A.; Baldi, P.; Rychnovsky, S. D.; Huang, L. Development of a Novel Cross-Linking Strategy for Fast and Accurate Identification of Cross-Linked Peptides of Protein Complexes. Molecular & Cellular Proteomics 2011, 10(1), M110.002170. https://doi.org/10.1074/mcp.M110.002212.

(53) Eaton, A. K.; Stewart, R. C. The Two Active Sites of Thermotoga Maritima CheA Dimers Bind ATP with Dramatically Different Affinities. Biochemistry 2009, 48 (27), 6412–6422. https://doi.org/10.1021/bi900474g.

(54) Hopkins, J. B.; Gillilan, R. E.; Skou, S. *BioXTAS RAW’.* Improvements to a Free Open-Source Program for Small-Angle X-Ray Scattering Data Reduction and Analysis. J Appl Crystallogr 2017, 50(5), 1545–1553. https://doi.org/10.1107/S1600576717011438.

(55) Franke, D.; Svergun, D. I. *DAMMIF, a* Program for Rapid *Ab-Initio* Shape Determination in Small-Angle Scattering. J Appl Crystallogr 2009, 42 (2), 342–346. https://doi.Org/10.1107/S0021889809000338.

(56) Schneidman-Duhovny, D.; Hammel, M.; Tainer, J. A.; Sali, A. Accurate SAXS Profile Computation and Its Assessment by Contrast Variation Experiments. Biophysical Journal 2013, 705 (4), 962–974. https://doi.Org/10.1016/j.bpj.2013.07.020.

(57) Schneidman-Duhovny, D.; Hammel, M.; Tainer, J. A.; Sali, A. FoXS, FoXSDock and MultiFoXS: Single-State and Multi-State Structural Modeling of Proteins and Their Complexes Based on SAXS Profiles. Nucleic Acids Res 2016, 44 (W1), W424–W429. https://doi.org/10.1093/nar/gkw389.

(58) Srivastava, M.; Georgieva, E. R.; Freed, J. H. A New Wavelet Denoising Method for Experimental Time-Domain Signals: Pulsed Dipolar Electron Spin Resonance. Journal of Physical Chemistry A 2017, 727 (12), 2452–2465. https://doi.org/10.1021/acs.jpca.7b00183.

(59) Srivastava, M.; Freed, J. H. Singular Value Decomposition Method to Determine Distance Distributions in Pulsed Dipolar Electron Spin Resonance. Journal of Physical Chemistry Letters 2017, 8(22), 5648–5655. https://doi.org/10.1021/acs.jpclett.7b02379.

(60) Bilwes, A. M.; Quezada, C. M.; Croal, L. R.; Crane, B. R.; Simon, M. I. Nucleotide Binding by the Histidine Kinase CheA. nature structural biology 2001, 8(4), 8.

(61) Park, S.-Y.; Borbat, P. P.; Gonzalez-Bonet, G.; Bhatnagar, J.; Pollard, A. M.; Freed, J. H.; Bilwes, A. M.; Crane, B. R. Reconstruction of the Chemotaxis Receptor-Kinase Assembly. Nat Struct Mol Biol 2006, 13(5), 400-07. https://doi.org/10.1038/nsmb1085.

(62) Stewart, R. C.; VanBruggen, R.; Ellefson, D. D.; Wolfe, A. J. TNP-ATP and TNP-ADP as Probes of the Nucleotide Binding Site of CheA, the Histidine Protein Kinase in the Chemotaxis Signal Transduction Pathway of *Escherichia Colit Biochemistry* 1998, 37 (35), 12269-12279. https://doi.org/10.1021/bi980970n.

(63) Hammel, M. Validation of Macromolecular Flexibility in Solution by Small-Angle X-Ray Scattering (SAXS). Eur Biophys J 2012, 41 (10), 789–799. https://doi.org/10.1007/s00249-012-0820-x.

(64) Putnam, C. D.; Hammel, M.; Hura, G. L.; Tainer, J. A. X-Ray Solution Scattering (SAXS) Combined with Crystallography and Computation: Defining Accurate Macromolecular Structures, Conformations and Assemblies in Solution. Quart. Rev. Biophys. 2007, 40 (3), 191—285. https://doi.org/10.1017/S0033583507004635.

(65) Rambo, R. P.; Tainer, J. A. Accurate Assessment of Mass, Models and Resolution by Small-Angle Scattering. Nature 2013, 496 (7446), 477–481. https://doi.org/10.1038/nature12070.

(66) Durand, D.; Vivès, C.; Cannella, D.; Pérez, J.; Pebay-Peyroula, E.; Vachette, P.; Fieschi, F. NADPH Oxidase Activator P67phox Behaves in Solution as a Multidomain Protein with Semi-Flexible Linkers. Journal of Structural Biology 2010, 769(1), 45–53. https://doi.Org/10.1016/j.jsb.2009.08.009.

(67) Swanson, R. V.; Bourret, R. B.; Simon, M. I. Intermolecular Complementation of the Kinase Activity of CheA. Mol Microbiol 1993, 8 (3), 435^41. https://doi.Org/10.1111/j. 1365-2958.1993.tb01588.x.

(68) Gräwe, A.; Stein, V. Linker Engineering in the Context of Synthetic Protein Switches and Sensors. Trends in Biotechnology 2021, 39 (7), 731–744. https://doi.org/l0.1016/j.tibtech.2020.11.007.

(69) Piñas, G. E.; Parkinson, J. S. Identification of a Kinase-Active CheA Conformation in Escherichia Coli Chemoreceptor Signaling Complexes. J Bactehol 2019, 207 (23). https://doi.org/10.1128/JB.00543-19.

(70) Hamel, D. J.; Zhou, H.; Starich, M. R.; Byrd, R. A.; Dahlquist, F. W. Chemical-Shift-Perturbation Mapping of the Phosphotransfer and Catalytic Domain Interaction in the Histidine Autokinase CheA from *Thermotoga Maritima ^Ť^ · \**. Biochemistry 2006, 45 (31), 9509–9517. https://doi.org/10.1021/bi060798k.

